# Genome-wide Screening Identifies Unique Host-Directed Drugs and Pro-viral Signalling Pathways for SARS-CoV-2

**DOI:** 10.1101/2025.10.15.682635

**Authors:** Juveriya Qamar Khan, Karthic Rajamanickam, Frederick S. Vizeacoumar, Mahrokh Balouchi, Yue Zhang, Hussain Elhasana, Megha Rohamare, He Dong, Kathleen Glover, Tristan Anderson-Woodsworth, Kalpana K. Bhanumathy, Jocelyne Lew, Anil Kumar, Franco J. Vizeacoumar, Darryl Falzarano, Joyce A. Wilson

## Abstract

SARS-CoV-2 is a positive-sense RNA virus and was responsible for the devastating COVID-19 pandemic. Although the current disease burden is less severe, there are limited treatment options, significant gaps in knowledge, and a looming threat of the emergence of variants and future pandemics. To address these challenges, we performed genome-wide CRISPR knockout screens in a novel human lung cell line NCI-H23^ACE2^, as well as in HEK293T^ACE2^ cells, with SARS-CoV-2 Wuhan virus, with the aim of identifying host-dependency factors that could predict effective antivirals. We identified four host-directed drugs, donepezil, dH-ergocristine, trametinib and sorafenib, that could potentially be repurposed to treat coronavirus infections. Three of the drugs inhibited SARS-CoV-2, HCoV-229E, and HCoV-OC43, suggesting they could be used as pan-coronavirus antivirals. We also confirmed that SARS-CoV-2 relies on the NRAS/Raf/MEK/ERK signaling pathway for its replication. Our study highlights the robustness and efficiency of a bilateral approach of gene silencing and antiviral screening to identify host-dependency factors and effective antivirals.

## Introduction

Coronaviruses are a diverse group of enveloped, single-stranded positive-sense RNA viruses, known to cause respiratory illnesses in humans and respiratory as well as intestinal tract infection in other mammals and avian species [1]. In humans, seasonal colds are commonly caused by human coronavirus (HCoV) −229E and NL63, belonging to the *alphacoronavirus* genus and HCoV-OC43, a *betacoronavirus* [2]. In 2002, Severe Acute Respiratory Syndrome Virus (SARS-CoV), a *betacoronavirus,* emerged causing severe respiratory illness, with a case fatality of ∼10%, [2, 3]. Ten years later, another highly pathogenic coronavirus, Middle East Respiratory Syndrome Virus (MERS-CoV), spreading from dromedary camels to humans, caused an outbreak in Saudi Arabia [2, 4]. Most recently, emerging in late 2019, SARS-CoV-2 (Severe Acute Respiratory Syndrome Virus-2), another *betacoronavirus*, quickly spread worldwide, resulting in the COVID-19 (Coronavirus Disease-2019) pandemic, facilitated by its high rate of transmissibility and ability to evolve rapidly. SARS-CoV-2 also shares ∼80% and ∼50% sequence identity with SARS-CoV and MERS-CoV respectively [1, 5]. It is believed that these coronaviruses emerged from bat reservoirs, and for SARS-CoV and MERS-CoV, masked palm civets and camels were identified as intermediary hosts, respectively [6]. Although pangolins and snakes have been speculated as intermediate hosts for SARS-CoV-2, the lack of surveillance data prior to the pandemic and poor evidence have made it challenging to implicate a single intermediate host for SARS-CoV-2 [7].

SARS-CoV-2 is primarily transmitted by respiratory droplets and aerosols expelled from infected individuals. Clinically, the virus can cause a wide range of symptoms, from mild flu-like symptoms to severe respiratory failure in individuals with co-morbidities [5]. SARS-CoV-2 replicates first in the epithelial cells in the respiratory tract and then makes its way to the alveolar epithelial cells in the lungs. This may trigger a strong immune response, resulting in a cytokine storm in pulmonary tissues through the hyperactivation of the immune system, which can cause acute to fatal respiratory distress [5, 8]. The prognosis of the elderly and people with underlying chronic conditions who are infected is relatively poor [5, 9]. In addition to acute effects, SARS-CoV-2 can cause long COVID or post-acute sequelae of COVID-19 (PASC), a multi-systemic condition that persists in patients after recovery from an acute severe COVID-19 infection [10, 11]. The symptoms of long COVID typically begin at about 4 weeks after infection, and can last weeks, months or even years. Research on long COVID is currently in the early stages and ongoing; however, several hypotheses have been suggested with the likelihood of multiple, overlapping causes, including persisting reservoirs of SARS-CoV-2 in tissues, impacts of the infection on the host microbiota, immune dysregulation with or without reactivation of underlying pathogens, microvascular blood clotting, autoimmunity, and dysfunctional signalling of the nervous system [10].

During infection, SARS-CoV-2 primarily enters cells by binding to the angiotensin-converting enzyme-2 (ACE2) cell surface protein receptor through its viral spike protein. Host proteases such as transmembrane protease, serine 2 (TMPRSS2) or cathepsin-L (CTSL) then cleave Spike to activate membrane fusion and entry into the cytoplasm [12]. Once the viral genome is released in the host cell cytoplasm, direct translation of the ORF1ab polyprotein takes place, using host cell machinery. The polyprotein is proteolytically processed into individual non-structural proteins that remodel host intracellular membranes to provide a protective and conducive environment for the replicase-transcriptase complex to initiate viral genomic replication and transcription of the subgenomic mRNAs [1, 13]. Viral genomic replication results in negative-sense genomic RNA copies and nested subgenomic RNAs, which function as templates for the generation of new positive-sense RNA genomes to be packaged, as well as the nested subgenomic mRNAs that are translated into structural and accessory proteins [13]. The structural proteins are then translocated into the endoplasmic reticulum (ER) membranes and transit through the ER-Golgi intermediate compartment (ERGIC), where interaction with the Nucleocapsid protein (N)-encapsidated genomic RNA facilitates the assembly of progeny virions and budding into the lumen of the secretory vesicular compartments [1, 13, 14]. The final stage of the virus life cycle involves exit of the virus from the infected cell through interaction with lysosomal trafficking pathways and exocytosis [1, 13, 14]. The complex life cycle within a host cell, necessitates that SARS-CoV-2 interacts with and hijacks several host factors and pathways during its infectious life cycle [1, 14].

While replicating, SARS-CoV-2, like all RNA viruses, continuously obtains and retains genetic mutations, thus giving rise to new genetic variants. During the stages of the pandemic, several new variants emerged and quickly became the predominant circulating variants. New variants typically had enhanced transmissibility, and immune escape that gave them advantages over previous variants. The Centers for Disease Control and Prevention (CDC) designates such rapidly evolving mutants with increased transmissibility and immune escape as variants-of-concern (VOCs) [15], and some had different disease severity. During the span of the pandemic, five SARS-CoV-2 variants became VOCs and were named Alpha, Beta, Gamma, Delta, and Omicron. Additional variants that never became predominant also emerged and were termed variants of interest (VOI) or variants under monitoring (VUM) by CDC and WHO. At present, sub-variants of Omicron, currently classified as VUMs, are known to be spreading through the human population, and their current global public health risk level is evaluated as low (https://www.who.int/activities/tracking-SARS-CoV-2-variants).

Emergence of variants makes it more challenging to keep using the same preventative or treatment options. The rapid development of several effective vaccines played a key role in controlling the disease burden of COVID-19, but none provide sterilizing immunity and virus spread continues. Treatment options for people who do become infected and have severe disease are limited. Direct acting antivirals such as remdesivir, molnupiravir and paxlovid (nirmatrelvir and ritonavir) have been approved by the United States Food and Drug Administration (FDA) for COVID-19 treatment. Remdesivir and molnupiravir are nucleoside analogs that inhibit the RNA dependent RNA polymerase (RdRp) enzyme, nirmatrelvir inhibits the SARS-CoV-2 main protease (Mpro), while ritonavir is a pharmacokinetic booster and inhibits hepatic metabolism of nirmatrelvir thus enhancing its plasma concentrations [16]. Direct acting antivirals, however, have reduced efficacy after the onset of severe disease, since virus replication is already reduced and the disease symptoms are a result of the hyperinflammatory response to infection, which can lead to multi-organ distress and Acute Respiratory Distress Syndrome (ARDS) [17]. For patients requiring oxygen supplementation, glucocorticoids such as dexamethasone were advised; however, it does not benefit patients who do not require oxygen [16]. Another popular treatment was the use of convalescent plasma from individuals recovered from past COVID-19 infection but is currently not recommended after randomized trials concluded it was not associated with reduction in severe COVID-19 progression [18]. Additionally, monoclonal antibodies that target the SARS-CoV-2 spike protein were used to treat several critical patients with COVID-19 under the FDA Emergency Use Authorizations (EUA). However, they are no longer recommended for use due to the new Omicron variants and subvariants not being susceptible to their treatment [16]. Another class of drugs that are used for COVID-19 are immunomodulatory drugs such as tocilizumab (interleukin-6 (IL-6) inhibitor), and baricitinib (Janna kinase (JAK) inhibitor) that have been approved by the FDA for use in hospitalized adults with COVID-19 (https://www.fda.gov/drugs/emergency-preparedness-drugs/coronavirus-covid-19-drugs [16]. Elevated levels of IL-6 were originally identified in association with SARS-CoV-related- and later in MERS-CoV related-severe respiratory distress [19, 20]. The known immunopathology of SARS-CoV and MERS-CoV resulted in an expedited approach to identify host-directed drugs such as tocilizumab, which can inhibit the strong cytokine response observed in COVID-19 patients [19]. Tocilizumab is indicated only for inpatients with oxygen requirements and is given with corticosteroids [16]. Thus, there is a strong requirement for newer and more effective treatment options for outpatients as well as hospitalized patients with COVID-19. Since coronaviruses hijack host factors and pathways, host-directed therapeutics can provide effective alternatives to traditional drugs, and since many coronaviruses hijack common pathways, host-directed therapeutics have the potential to treat a broad-spectrum of viruses and could enhance our preparation for future coronavirus outbreaks.

One strategy for developing antiviral therapeutics is to repurpose existing drugs that may have antiviral activity. To identify drugs with potential antiviral activity, we used a genome-wide CRISPR knockout (KO) screen to identify host factors and pathways that are used by SARS-CoV-2, termed host dependency factors, and then tested drugs that target these host factors for their antiviral activity. Several genome-wide CRISPR KO screens to identify SARS-CoV-2 host dependency factors have been performed so far [21–29]; however, ours is unique in that it was performed using a novel human lung cell line. CRISPR screens rely on a library of guide RNAs (gRNAs) that target all genes in the human genome to generate a population of cells in which theoretically one gene has been knocked out per cell. Then these cells are infected by the “virus-to-be-tested”, for lethal rounds of infection. Cells that survive the infection potentially have a knockout of a gene that is required for efficient virus replication, and thus the screen relies on live-dead selection of virus-susceptible and resistant cells. Thus, cells that are highly susceptible to virus-induced cell death is a major criterion for the choice of a cell line. Cell lines commonly used in CRISPR screens for SARS-CoV-2 include Calu3, and Vero, or cell lines transduced to overexpress ACE2 and/or TMPRSS2 to increase virus susceptibility such as A549, Huh7/7.5, CaCo-2 and 293T. Of these, Calu3 and A549 are the only representative lung cell lines used for CRISPR screens so far [21–29]. Unique to our study, we used a novel lung adenocarcinoma cell line NCI-H23^ACE2^, to perform genome-wide CRISPR KO screens for SARS-CoV-2. NCI-H23^ACE2^ is highly susceptible to SARS-CoV-2 infection and shows robust virus-induced cell cytopathic effect (CPE), with up to 99% cell death [30]. We used this cell line to highlight lung-relevant gene knockouts to identify important cellular pathways and functions used by SARS-CoV-2, thus complementing other studies and at the same time providing unique findings. Based on the CRISPR screen host dependency factors, our study identified four potential antiviral drugs, donepezil, dihydroergocristine (dH-ergocristine), trametinib and sorafenib, that inhibit SARS-CoV-2, HCoV-229E and HCoV-OC43 replication. Based on these drug targets in addition to siRNA knockdown validations, we also confirmed a pro-viral role of the NRAS/Raf/MEK/ERK pathway. We were also able to identify several other potential pro-viral host factors, including several ribonucleoprotein complex genes, including RPL3, RPL18A and APOBEC3F; ciliogenesis associated gene BBS1; and KAT5, a lysine acetyltransferase.

## Results

### CRISPR KO screens for SARS-CoV-2 identified unique gene hits in HEK293T^ACE2^ and NCI-H23^ACE2^ cells

To identify important host dependency factors required by SARS-CoV-2, we performed CRISPR KO screens in two susceptible cell lines, HEK293T^ACE2^ and NCI-H23^ACE2^. The GeCKO gRNA library B was used and has three gRNAs targeting each of 19,050 genes, along with 1000 non-targeting control gRNAs. Cas9 expressing HEK293T^ACE2^ cells were transduced with gRNA library and then infected with SARS-CoV-2 Canada/ON/VIDO-01-2020, a lineage B Wuhan1 isolate at an MOI of 0.3, and total cellular DNA was collected when we observed ∼80% virus-induced CPE. To increase the stringency of our screening conditions, resistant HEK293T^ACE2^ cells were reinfected with SARS-CoV-2 two more times at 48-hour intervals. At day 6 post-infection, genomic DNA from surviving cells was extracted, amplified, and sequenced to identify transduced gRNAs. To identify lung-specific host dependency factors, we also performed the screen with an adenocarcinoma cell line, NCI-H23^ACE2^ that is highly susceptible to SARS-CoV-2 infection and exhibits ∼99% virus-induced CPE with SARS-CoV-2 Wuhan VIDO-01 virus at MOI=0.5, 72 hours post-infection [30]. Upon infection with SARS-CoV-2 VIDO-01 at MOI=0.1, virus resistant NCI-H23^ACE2^ cells were collected when we observed ∼95% CPE, at 48-72 hours post-infection and genomic DNA was extracted to be processed for amplifying the guide sequences by PCR and subsequent next-generation sequencing (NGS).

Using MAGeCK analysis, hits were ranked according to their enrichment score (z-score) with a cut-off of 3.5 and p <0.05. The screen done in NCI-H23^ACE2^ gave a list of 430 potential pro-viral host genes (Figure 1a) (Supplementary table S1) and the HEK293T^ACE2^ screen resulted in 296 potential pro-viral genes (Figure 1c) (Supplementary table S2). The top ten hits from each cell line are depicted in the respective volcano plots (Figure 1a, c). Gene Ontology (GO) enrichment analysis found several host cellular processes important for SARS-CoV-2 in both NCI-H23^ACE2^ and HEK293T^ACE2^ (Figure 1b, d). Some of the enriched GO annotations indicated pathways important for virus infection, including, intracellular signal transduction, protein phosphorylation, intracellular transport, protein containing complex assembly, cytoskeleton organization, cellular components of vesicle membranes, and endosome membrane (Figure 1b, d) (Supplementary table S3, S4).

**Figure 1.**
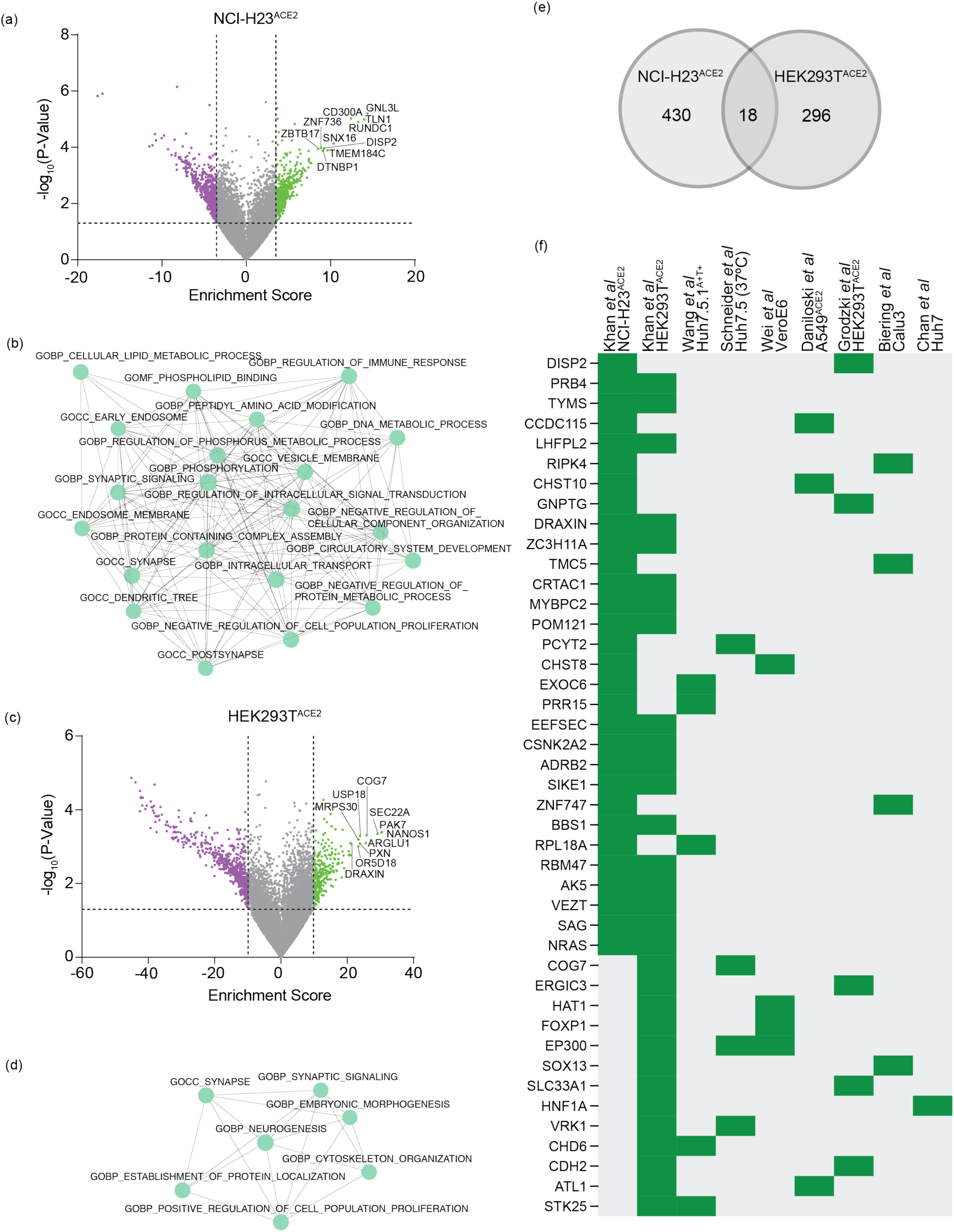
CRISPR KO screens of NCI-H23^ACE2^ and HEK293T^ACE2^, reveal SARS-CoV-2 host dependency factors and pathways. (a) Analysis of CRISPR KO screen for SARS-CoV-2 Wuhan VIDO-01, in NCI-H23^ACE2^ cells reveal enriched genes that are represented as a volcano plot; wherein the SARS-CoV-2 host-dependency genes hits, represented as green dots were ranked according to their enrichment score (x-axis), with a cut-off of 3.5 and with p<0.05 (y-axis). The top 10 ranking hits (GNL3L, TLN1, RUNDC1, CD300A, SNX16, TMEM184C, DISP2, DTNBP1, ZNF736, ZBTB17) are labelled in the volcano plot. The screen represents data from three independent experiments. (b) Gene ontology (GO) analysis of host-dependency hits from the NCI-H23^ACE2^ screen reveal enriched cellular networks important for SARS-CoV-2 life cycle, annotated as GOBP (gene ontology biological process), GOMF (gene ontology molecular function) and GOCC (gene ontology cellular component). (c) Analysis of CRISPR KO screen for SARS-CoV-2 Wuhan VIDO-01, in HEK293T^ACE2^ cells reveal enriched genes that are represented as a volcano plot; wherein the SARS-CoV-2 host-dependency genes hits, represented as green dots were ranked according to their enrichment score (x-axis), with a cut-off of 3.5 and p <0.05 (y-axis). The top 10 ranking hits (PAK7, NANOS1, SEC22A, COG7, ARGLU1, USP18, PXN, MRPS30, OR5D18 and DRAXIN) are labelled in the volcano plot. (d) GO analysis of host-dependency hits from the HEK293T^ACE2^ screen reveal enriched cellular networks important for SARS-CoV-2 life cycle, annotated as GOBP, and GOCC. (e) We identified a total of 430 gene hits from NCI-H23^ACE2^ screen and 296 gene hits from HEK293T^ACE2^ screen, with 18 overlapping putative hits from both our screens, represented as a Venn diagram. (f) Meta-analysis of our screens and with other CRISPR KO studies reveal overlapping gene hits, represented as green blocks; with the genes labelled on the left, and the study and cell line used for the CRISPR screen labelled above the graph [21–29]. The genes are arranged according to the p-value calculated for NCI-H23^ACE2^ and HEK293T^ACE2^ screen in our study.

We identified 18 putative host dependency factors common to both our screens in HEK293T^ACE2^ and NCI-H23^ACE2^ cells (Figure 1e, f), including, ADRB2, BBS1, CSNK2A2, MYBPC2, NRAS, PRB4, SAG, TYMS, ZC3H11A, POM121, LHFPL2, AK5, RBM47, CRTAC1, VEZT, EEFSEC, SIKE1 and DRAXIN. Furthermore, meta-analyses of our screens in NCI-H23^ACE2^ and HEK293T^ACE2^ cell lines, with CRISPR KO screens done by other groups [21–29] identified 12 and 13 common putative host dependency factors, respectively, as shown in Figure 1f, including COG7, EXCO6 and RPL18A. Owing to variables such as different cell lines, the CRISPR guide RNA libraries, scoring methods, and assay conditions, our study also resulted in several unique hits, such as KAT5, HTR3E, and WNT4.

### Unique host-directed drugs that can be repurposed for SARS-CoV-2 were identified by our antiviral screening approach

Several of the genes in our screens can be targeted by FDA-approved or experimental drugs (https://www.cancerrxgene.org/, https://www.genecards.org/). We selected 21 such drugs that target genes identified as pro-viral in our CRISPR screens (Supplementary table S5). To assess the antiviral activity of each of the drugs against SARS-CoV-2, A549^ACE2^ cells were used. A549^ACE2^ cells have been used previously for SARS-CoV-2 antiviral assays [24, 31, 32] and allows for validation of our screen in another cell line. The cells were pre-treated with drugs at four concentrations, infected with SARS-CoV-2 Wuhan NLuc virus and incubated with the drug for another 48 hours before assessing the virus replication activity by luciferase assay. The antiviral activity of 22 drugs tested is represented as a heatmap (Figure 2a), wherein remdesivir was used as a positive control, and is seen to inhibit viral activity strongly at the highest concentrations (Figure 2a). All the drugs were used at concentrations that maintained at least 70% cell viability (Figure S2). We identified that Donepezil, dihydroergocristine (dH-ergocristine), and Trametinib inhibited virus replication based on robust inhibition of viral luciferase activity (Figure 2a). Donepezil and dH-ergocristine are novel to this study. Sorafenib also inhibited, but with relatively less potency (Figure 2a).

**Figure 2.**
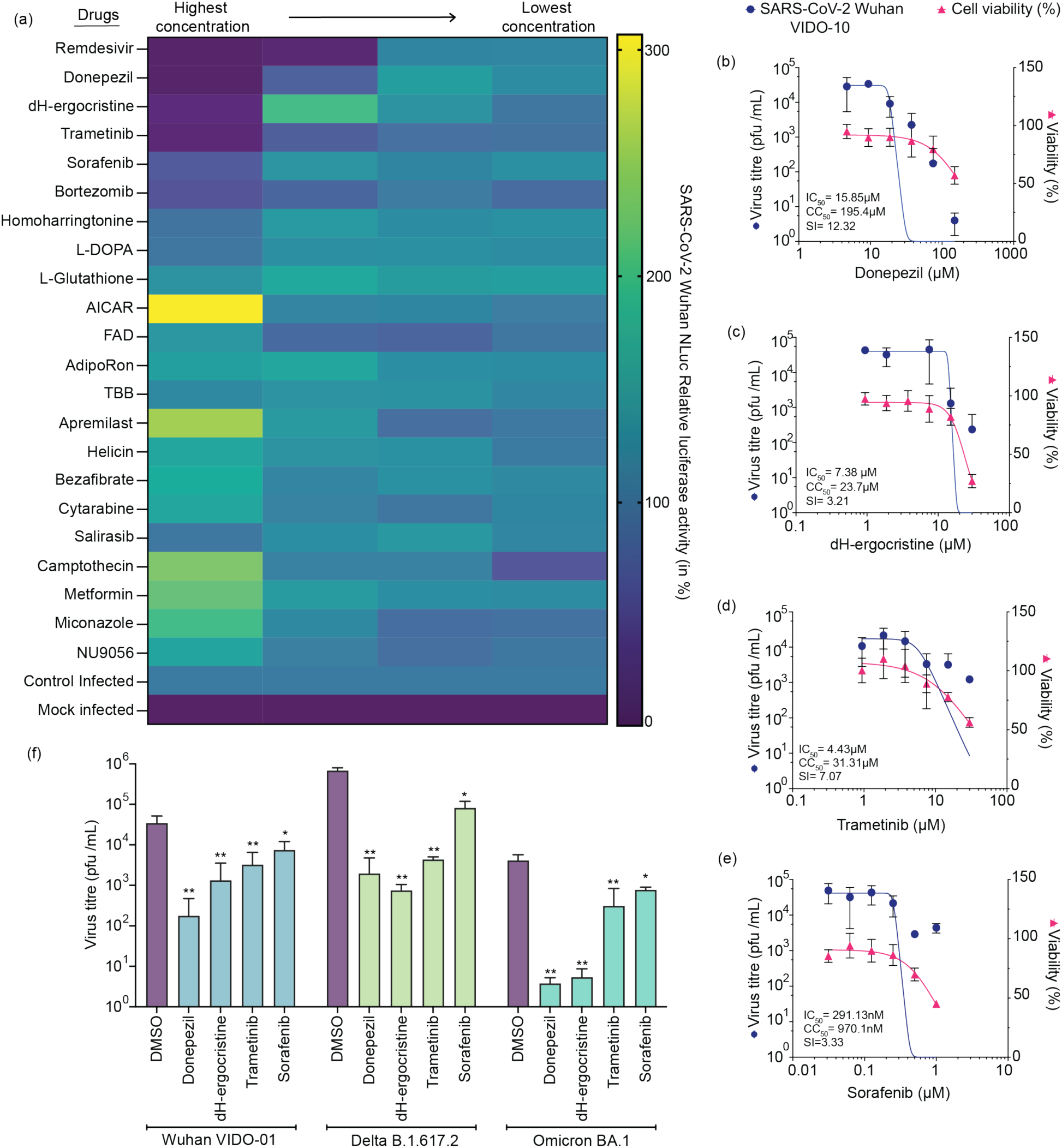
Screening of host-directed drugs identified unique drugs that inhibit SARS-CoV-2 *in vitro*. (a) FDA-approved or experimental drugs were screened against SARS-CoV-2 Wuhan NLuc virus in A549^ACE2^ cells to test their antiviral activity. Briefly, cells were pre-treated at four concentrations of each drug (Supplementary table S5) followed by virus infection in the presence of the drug, and further incubation with the drug for 48 hours. Virus activity was assessed by luciferase assay, represented in the heatmap as relative luciferase activity (in %), normalized to DMSO control infected cells (represented at 100% infection as blue bars). Mock infected cells are represented at 0% infection as purple bars. The scale of luciferase activity is indicated on the right. The corresponding cell viability for each drug is shown in Supplementary Figure S2. (b-f) From the antiviral screen, four drugs that inhibited SARS-CoV-2 luciferase at the highest concentrations, were further tested against wild-type SARS-CoV-2 Wuhan VIDO-01 virus to confirm inhibition of virus titres, in A549^ACE2^ cells. The drug-dose response curves were generated using non-linear regression analysis, for (b) donepezil, (c) dH-ergocristine, (d) trametinib and (e) sorafenib, with the x-axis indicating the concentration (in µM) of each drug. The left y-axis depicts virus titers (in plaque forming units (pfu)/mL) in drug-treated cells (blue line graph), and the right y-axis depicts relative cell viability (in %) in uninfected drug-treated cells, normalized to DMSO-treated cells, (pink line graph). The 50% inhibitory concentration (IC_50_), 50% cytotoxicity concentration (CC_50_) and selectivity index (SI) (ratio of CC_50_/ IC_50_) are labelled in each graph. (f) The four shortlisted drugs, donepezil (75 µM), dH-ergocristine (15 µM), trametinib (15 µM), and sorafenib (0.5 µM), were further confirmed to reduce virus titres of SARS-CoV-2 variants, Delta B.1.617.2 and Omicron BA.1 at the indicated concentrations. The y-axis represents virus titers (in pfu/mL) when treated with DMSO (purple bar) or drugs (shades of green) as labelled on the x-axis and infected with wild-type SARS-CoV-2 Wuhan VIDO-01, Delta B.1.617.2 or Omicron BA.1. The data represent an average of at least three independent experiments and error bars represent the standard deviation. Statistical significance was determined using two-way ANOVA and compared to DMSO control, for each variant; where ns p>0.1234, * p<0.0332, ** p<0.0021, *** p<0.0002, **** p<0.0001.

Donepezil has been reported to target lysine acetyltransferase 5 (KAT5), a protein identified in our screen, and is a drug initially developed for treatment of Alzheimer’s disease [33]. DH-ergocristine targets 5-Hydroxytryptamine Receptor 3E (HTR3E), a receptor subunit of 5-Hydroxytryptamine or serotonin, identified as a host dependency factor in our CRISPR screen. dH-ergocristine is an FDA-approved drug used for the treatment of cognitive disorders such as dementia and is a serotonin receptor antagonist [34, 35]. Trametinib inhibits MEK1/2 (mitogen-activated extracellular signal-regulated kinase) and was selected based on its targeting of NRAS pathway. NRAS (the neuroblastoma RAS viral [v-ras] oncogene homolog) is a gene identified in our CRISPR screen and initiates the NRAS/Raf/MEK/ERK pathway [36]. In the same pathway, sorafenib inhibits serine/threonine kinases including Raf (rapidly accelerated fibrosarcoma) and was chosen based on its inhibition of MAPK1 (mitogen-activated protein kinase), a gene identified in our CRISPR screen, through inhibition of Raf upstream of the MAPK1 activation. MAPK1 is also known as ERK2 (Extracellular Signal-Regulated Kinase 2) and is another component of the NRAS pathway [37]. The antiviral effects of donepezil, dH-ergocristine, trametinib, and sorafenib were further confirmed in dose response assays to assess reductions in viral titers of SARS-CoV-2 Wuhan VIDO-01 (Figure 2b) and to calculate the 50% inhibitory concentration (IC_50_), 50% cytotoxicity concentration (CC_50_) and selectivity index (SI). Donepezil inhibited SARS-CoV-2 VIDO-01 with an IC_50_ of 15.85 µM and SI of 12 (Figure 2b); whereas, dH-ergocristine inhibited SARS-CoV-2 with an IC_50_ of 7.38 µM and an SI 3.21 (Figure 2c). Trametinib and sorafenib treatment also reduced SARS-CoV-2 titers and the IC_50_ was calculated as 4.43 µM and 0.29 µM resulting in an SI of 7.07 and 3.33 respectively (Figure 2d, e). Bortezomib, a proteasomal inhibitor, showed some inhibition of virus luciferase activity in our drug screen (Figure 2a) and in other studies [38, 39]. However, in our study it did not inhibit SARS-CoV-2 viral titers significantly at a concentration non-toxic for the cells so was not investigated further (data not shown). This could potentially indicate an inhibition of luciferase activity by bortezomib, without affecting virus activity.

### Host-directed drugs identified in our screen have pan-anti-coronaviral activity

Donepezil, dH-ergocristine, trametinib and sorafenib were further tested for their antiviral activity against other SARS-CoV-2 variants Delta B.1.617.2 and Omicron BA.1, at concentrations 2-3 fold higher than the IC_50_ to ensure significant inhibition of virus titers, while maintaining the cell viability at >70% (Figure 2b-f) (Supplementary table S5). All four drugs significantly reduced viral titers of SARS-CoV-2 Delta B.1.617.2 and Omicron BA.1 variants (Figure 2f). In all, our study has successfully identified novel drugs that can be repurposed against SARS-CoV-2 and that are effective against multiple SARS-CoV-2 variants.

Donepezil, dH-ergocristine, trametinib and sorafenib were also tested against common cold coronaviruses, HCoV-229E and HCoV-OC43, to examine their pan-coronaviral inhibitory capacity. The antiviral assay was carried out as described above in Huh-7 cells for HCoV-229E and in MRC-5 cells for HCoV-OC43. Donepezil was effective against HCoV-229E and HCoV-OC43 at IC_50_ 12.5µM and 23.21µM, respectively (Figure 3a, e), whereas dH-ergocristine was effective at IC_50_ of 1.12 µM and 2.29 µM, respectively, for each virus (Figure 3b, f). Furthermore, trametinib also significantly inhibited HCoV-229E and HCoV-OC43 with an IC_50_ of 1.35µM and 9.7µM, respectively (Figure 3c, g). That the SI for each of these drugs against HCoV-229E and HCoV-OC43 is >4.7, indicates that donepezil, dH-ergocristine, and trametinib have a pan-coronaviral inhibitory effect. Sorafenib, although inhibitory against HCoV-229E with an IC_50_ of 510nM and SI of 7.3, did not significantly inhibit HCoV-OC43 titers (Figure 3d, h). Figure 3i summarizes the SI and thus the corresponding antiviral activity, of each of the four drugs against SARS-CoV-2, HCoV-229E and HCoV-OC43.

**Figure 3.**
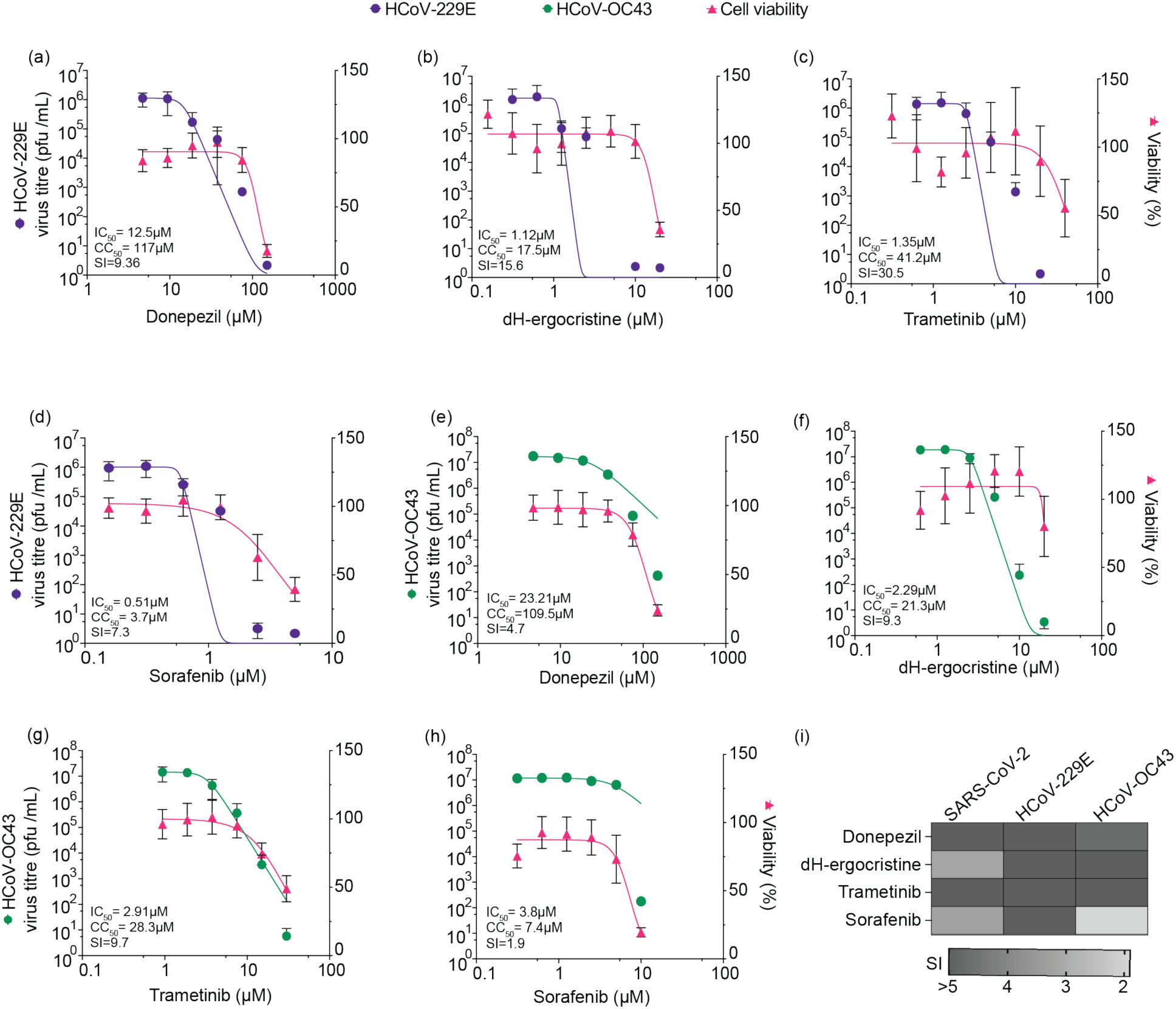
Host-directed drugs identified in our screen can be repurposed as anti-pan-coronaviral drugs. (a) Donepezil, (b) dH-ergocristine, (c) trametinib and (d) sorafenib were tested against human coronavirus HCoV-229E in Huh-7 cells. The drug dose response curves were generated using non-linear regression analysis, with concentrations (in µM) of each drug along the x-axis; the left y-axis depicts virus titers (in pfu/mL) in drug-treated cells corresponding to the purple line graph, and the right y-axis depicts relative cell viability (in %) in uninfected drug-treated cells, normalized to DMSO-treated cells, corresponding to the pink line graph. The IC_50_, CC_50_ and SI are labelled in each graph. The data represent an average of at least three independent experiments and error bars represent the standard deviation. (e) Donepezil, (f) dH-ergocristine, (g) trametinib and (h) sorafenib were further tested against human coronavirus HCoV-OC43 in MRC-5 cells. The drug-dose responses curves were generated using non-linear regression analysis, with concentrations (in µM) of each drug along the x-axis; the left y-axis depicts virus titers (in pfu/mL) in drug-treated cells corresponding to the green line graph, and the right y-axis depicts relative cell viability (in %) in uninfected drug-treated cells, normalized to DMSO-treated cells, corresponding to the pink line graph. The IC_50_, CC_50_ and SI are labelled in each graph. The data represent an average of at least three independent experiments and error bars represent the standard deviation. (i) A heatmap summarizes SI and consequently the effective antiviral activity of donepezil, dH-ergocristine, trametinib and sorafenib against SARS-CoV-2, HCoV-229E and HCoV-OC43. The graph illustrates that donepezil and trametinib have the strongest antiviral activity against all three coronaviruses tested, with an SI of >4 indicated by darker grey shades. dH-ergocristine has an SI of >5 for the common cold coronaviruses and >3 for SARS-CoV-2. Sorafenib was effective against SARS-CoV-2 and HCoV-229E with an SI of >3 but not HCoV-OC43.

### Combinations of the host-directed drugs identified in our screen can inhibit SARS-CoV-2 in a more than additive manner and show inhibition in two human lung cell lines

Enhanced inhibitory effect by synergism between two drugs can allow for lower doses of each to be used. This could reduce undesirable side effects and avoid the evolution of resistant viruses. To test for synergism between the four drugs we tested two-at-a-time combinations at their IC_50_ concentrations against wild type viruses-SARS-CoV-2 Wuhan VIDO-01, Delta B.1.617.2 and Omicron BA.1 variants in A549^ACE2^ cells (Figure 4). The drugs were first confirmed to show minimal cell toxicity when combined (Figure 4a), and viral titers in cell supernatants were then tested 48 hours post-infection. Using the Bliss Independence Model for drug combination, synergistic inhibition was defined by higher cumulative inhibition of two drugs together, compared to the single drug treatment or simply their additive inhibition (evaluated in Supplementary table S6) [40]. DH-ergocristine in combination with trametinib showed higher inhibition compared to expected additive drug inhibition, against SARS-CoV-2 Wuhan VIDO-01, and Omicron BA.1 potentially suggesting synergistic inhibition (Figure 4b, c, d, Supplementary table S6). Similarly, dH-ergocristine further shows greater than additive inhibition with sorafenib against Omicron BA.1 and Delta B.1.617.2 variants, as well as a combination of trametinib and sorafenib against all three variants (Figure 4b, c, d, Supplementary table S6). This indicates that dH-ergocristine, sorafenib, and trametinib are strong contenders for combination therapy against SARS-CoV-2.

**Figure 4.**
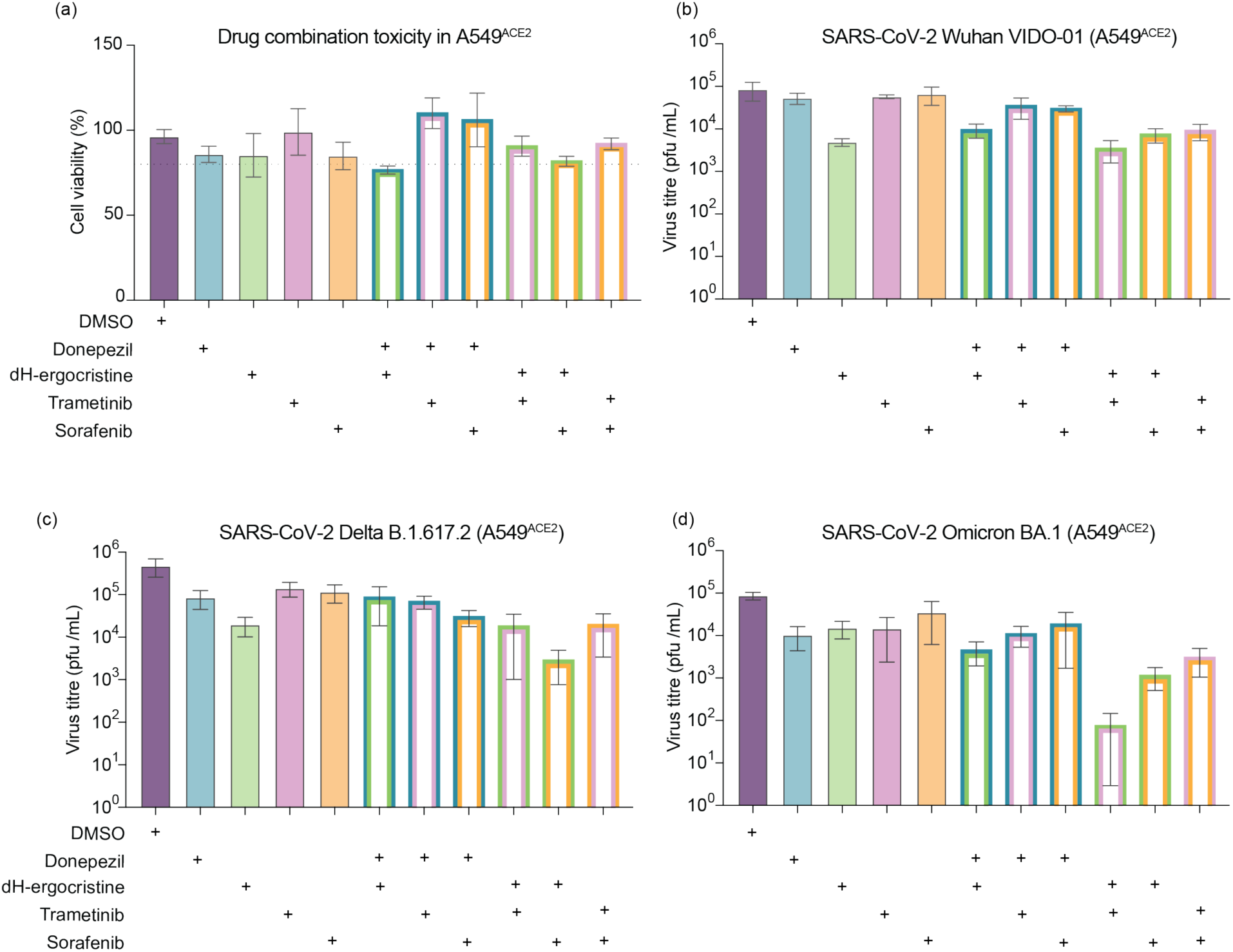
Specific combinations of effective host-directed antiviral drugs identified in our study have more than just additive inhibition against SARS-CoV-2. The four shortlisted drugs, donepezil, dH-ergocristine, trametinib and sorafenib, were tested in combinations of two in A549^ACE2^ cells and, (a) were first confirmed to be non-toxic for the cells using an ATP based cell viability assay. The drug combinations were performed with (b) SARS-CoV-2 Wuhan VIDO-01, (c) Delta B.1.617.2, and (d) Omicron BA.1. The y-axis represents virus titers (in pfu/mL) when treated with DMSO (purple bar) or drugs (blue-donepezil, green-dH-ergocristine, pink-Trametinib, orange-Sorafenib) as labelled on the x-axis with (+). For the drug combination assays, virus titers are represented as double bordered bar graphs and the respective drug combinations are indicated on the x-axis with (+). The data represent an average of at least three independent experiments and error bars represent the standard deviation.

Next, to confirm that the antiviral effect of the drugs is not cell line specific, we tested their efficacy against SARS-CoV-2 Wuhan and Delta in Calu3 cells, using reporter NLuc viruses. The IC_50_ concentrations, as calculated for A549^ACE2^ cells, was included in the range of concentrations we tested in Calu3 cells (Figure 5, Supplementary table S5). While maintaining minimal cell toxicity (Figure 5a), dH-ergocristine and trametinib showed inhibition of Wuhan NLuc (Figure 5b) and Delta NLuc (Figure 5c), viruses in a dose dependent manner. This indicates that dH-ergocristine and trametinib, are effective against SARS-CoV-2 in multiple cell lines. We further tested these drugs in combinations in Calu3 cells at concentrations showing no/minimal cell toxicity (Figure 5d, Supplementary table S5). Similar to what we saw in A549^ACE2^ cells, the combinations of trametinib and dH-ergocristine, as well as sorafenib and dH-ergocristine, displayed Bliss synergistic inhibition against SARS-CoV-2 Delta and Wuhan NLuc viruses (Figure 5e, 5f, Supplementary table S6). Additionally, donepezil and dH-ergocristine also showed higher than additive inhibition against both NLuc variants, whereas, trametinib and sorafenib showed Bliss synergy against Delta NLuc virus (Figure 5e, 5f, Supplementary table S6). This corroborated our finding on synergistic inhibition with some of the drug combinations in two independent cell lines.

**Figure 5.**
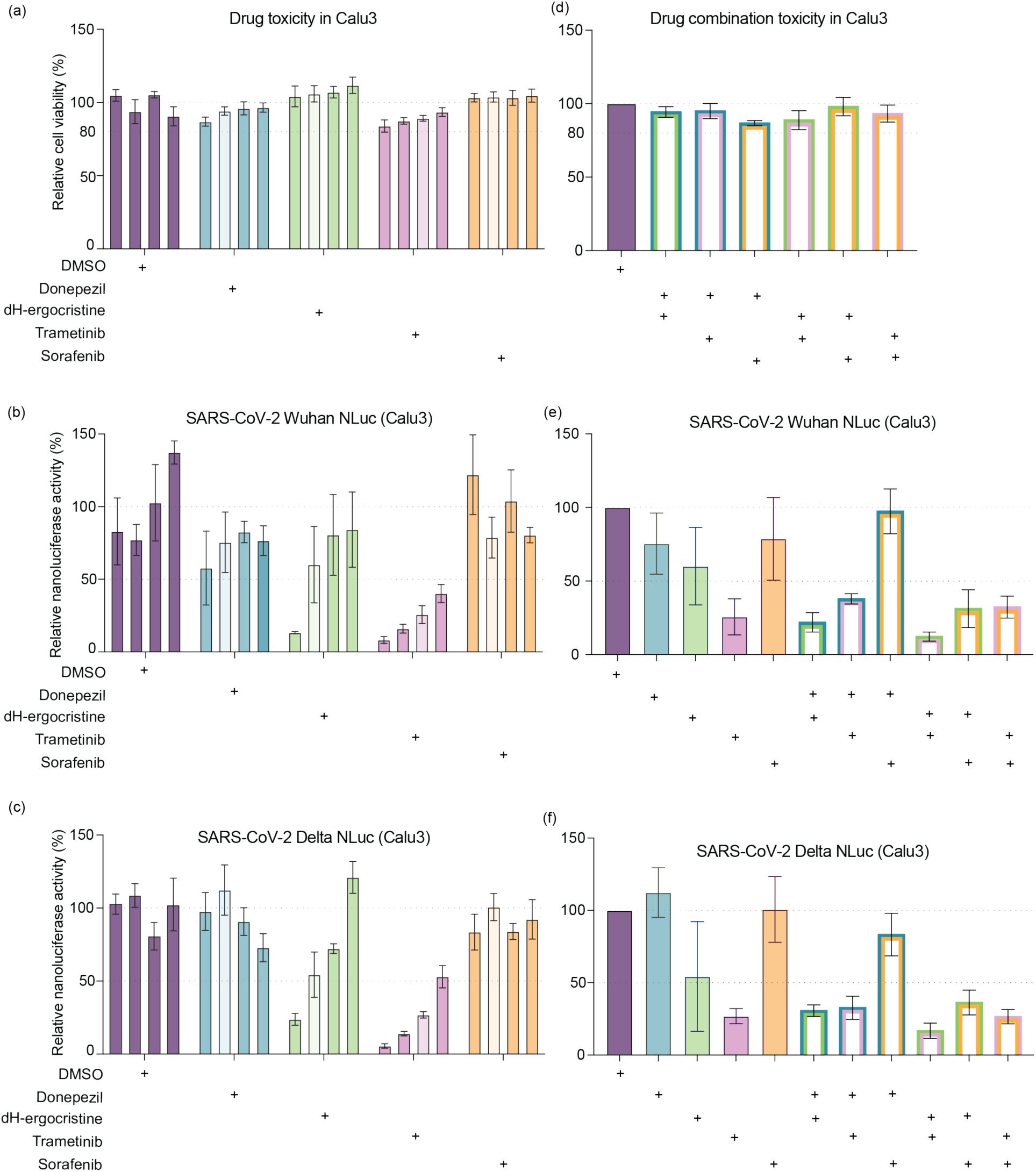
DH-ergocristine and trametinib inhibit SARS-CoV-2 Wuhan and Delta NLuc viruses in another human lung cell line, Calu3, and suggest higher inhibition in combination. (a) Treatment of donepezil (blue bars), dH-ergocristine (green bars), trametinib (pink bars), and sorafenib (orange bars) at four concentrations, including the IC_50_ values as calculated in A549^ACE2^cells (indicated in lighter shades), did not adversely affect cell viability in Calu3 cells as compared to DMSO treated cells (purple bars). The four drugs at four concentrations were tested against (b) SARS-CoV-2 Wuhan NLuc and (c) SARS-CoV-2 Delta NLuc. (d) Treatment of drugs in combinations (double bordered bars), as indicated on the x-axis (+: drug treated with), was also confirmed to not adversely affect cell viability in Calu3 cells as compared to DMSO treated cells (purple bars). The drug combinations were tested against (e) SARS-CoV-2 Wuhan NLuc and (f) SARS-CoV-2 Delta NLuc viruses. The data represent an average of at least three independent experiments and error bars represent the standard deviation.

### Secondary siRNA knockdown screen with a SARS-CoV-2 reporter virus further short-listed pro-viral gene hits

Apart from potential antivirals, the CRISPR KO screen gave us a list of potential pro-viral gene hits. To validate these hits, we performed a secondary siRNA knockdown screen in NCI-H23^ACE2^ cells to assess the impact on virus replication. We chose 165 genes to be tested, including the top 10 hits from both the NCI-H23^ACE2^ and HEK293T^ACE2^ screens (Figure 1a, c), 18 genes overlapping both the screens (Figure 1f), 20 gene hits that we targeted with available drugs (Figure 2), and the rest were chosen because they were lung specific genes or pathways required by other viruses. For these selected genes, we tested the impact on virus replication after transfection with a panel of four pooled siRNAs per gene (Supplementary table S7). To confirm efficient siRNA transfection and knockdown in NCI-H23^ACE2^ cells, we optimized siRNA knockdowns using an siRNA (siDUSP11), which was previously shown to achieve robust knockdown of DUSP11 (Dual Specificity Phosphatase 11 that interacts with RNA/RNP Complex) [41]. With the successful knockdown of DUSP11 confirmed by Western blot assay (Figure S3), we proceeded with the siRNA transfection screening. Briefly, siRNAs were transfected into NCI-H23^ACE2^ cells, which were then incubated for 48 hours to allow for knockdown of the respective proteins. Following this, the cells were infected with SARS-CoV-2 Wuhan NLuc virus for rapid assessment of replication efficiency. 24 hours post-infection, virus activity was assessed by luciferase assay, which was normalized to cells transfected with a non-targeting siRNA-siControl. siCTSL, an siRNA targeting a known SARS-CoV-2 host dependency factor CTSL, was used as positive control and we observed reduced virus luciferase activity after siRNA knockdown (Figure 6a). To determine if the siRNA transfection had adverse effects on essential cellular functions, transfected cells were left uninfected in parallel and were assayed for cell viability. From the 165 genes tested (Supplementary table S7), we identified seven siRNA pools that decreased virus luciferase values by more than 50% (RPL18A, APOBEC3F, RPL3, TBC1D15, IL36A, PSMA2, DISP2) (Figure 6a) (Supplementary table S7). We also identified 55 gene-targeting siRNAs that reduced virus luciferase activity significantly (Figure 6a). Our siRNA screen identified eight common putative host dependency hits from our CRISPR KO screens in both cell lines, and other CRISPR KO screens (Figure 1f, 6a), supporting the robustness of our screens. These include DISP2 [26], RPL18A [21], and BBS1, MYBPC2, POM121, CRTAC1, CSNK2A2 and NRAS (Figure 1f, 6a).

**Figure 6.**
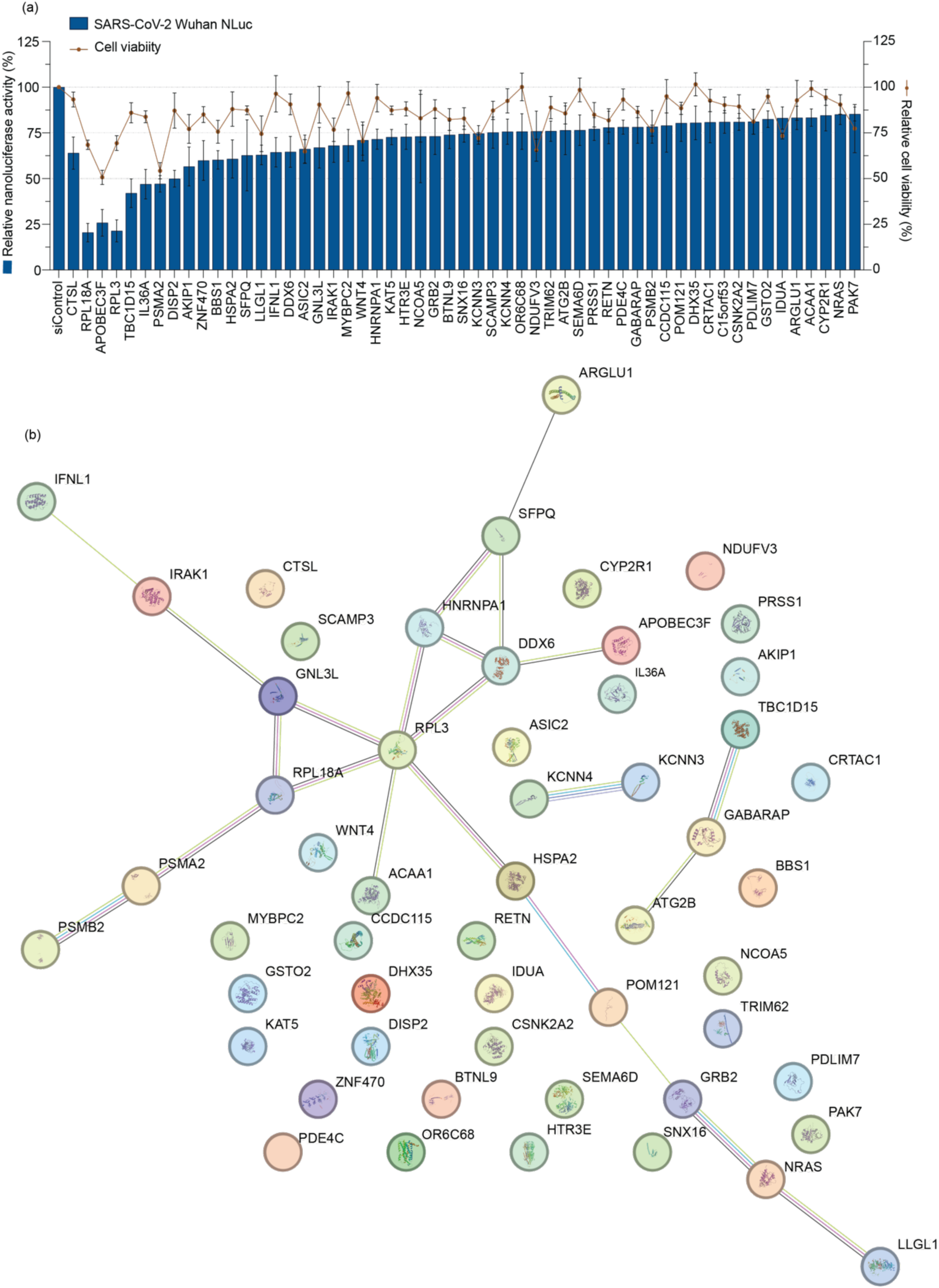
Secondary siRNA knockdown screen in NCI-H23^ACE2^ cells identifies important SARS-CoV-2 host dependency factors. (a) To confirm the potential pro-viral nature of the CRISPR KO hits, a secondary screen with 165 genes was carried out in NCI-H23^ACE2^ cells. Pools of four siRNAs against each gene was transfected in cells followed by infection with SARS-CoV-2 Wuhan NLuc virus. NLuc activity was assessed 24 hours post infection. The top 55 genes, indicated on the x-axis, reduced virus activity post-siRNA knockdown significantly, the rest of the genes did not (Supplementary table 7). Virus activity, assessed by a luciferase assay, is represented as blue bars on the graph corresponding with the left y-axis denoting relative luciferase activity (in %), normalized to infection with a non-targeting siRNA, siControl. In parallel, cell viability was assessed in uninfected siRNA transfected cells and is depicted respectively for each gene as a red line graph corresponding with the right y-axis denoting relative cell viability (in %), normalized to cell viability in siControl transfected cells. The data represent an average of at least three independent experiments and error bars represent the standard deviation. Statistical significance was determined using two-way ANOVA and compared to siControl; and the top 55 significant gene hits are shown in the figure. (b) STRING network analysis of the top 55 genes from the secondary screen, resulted in a comprehensive protein-protein association network, with proteins represented as colored circles or nodes and the connecting lines indicating possible protein interactions.

Of the 55 genes to which siRNA targeting showed an impact on SARS-CoV-2, we used the STRING-db v12 network analysis tool to identify common functionalities and networks. This resulted in a protein-protein association network as depicted in Figure 6b. Several proteins were functionally annotated to ribonucleoprotein (RNP) complexes and RNA metabolism, including GNL3L, RPL18A, RPL3, APOBEC3F, HNRNPA1, SFPQ, DDX6, and DHX35 (Figure 6b). The top three pro-viral genes identified in our siRNA screen were ribonucleoproteins, RPL18A, APOBEC3F and RPL3 that showed >75% reduction in infection, albeit with a decreased cell viability (Figure 6b), suggesting that ribonucleoprotein complexes play a role in SARS-CoV-2 replication and indicate an intricate viral RNA-host protein interactome requirement for the SARS-CoV-2 life cycle. Other pathways identified to be pro-viral included mitophagy (TBC1D15, NRAS, GABARAP, CSNK2A2), P-body (DDX6, APOBEC3F, PSMA2), proteins with calmodulin binding (KCNN3, KCNN4), cytokine signalling in immune responses (IL36A, TRIM62, IRAK1, IFNL1, POM121, GRB2, NRAS, PSMA2, PSMB2) and several components of intracellular non-membrane-bounded organelles (GRB2, MYBPC2, NCOA5, BBS1, LLGL1, APOBEC3F, DDX6, SFPQ, RPL3, RPL18A, GNL3L, IRAK1, KAT5, PSMA2. HSPA2, BBS1, LLGL1) implying the importance of these structures and pathways in SARS-CoV-2 lifecycle.

### KAT5, HTR3E, NRAS, and GNL3L act as SARS-CoV-2 host dependency factors

Both of our drug and siRNA screens indicated that KAT5 (Lysine acetyltransferase 5), HTR3E (5-Hydroxytryptamine (Serotonin) Receptor 3 family member E), NRAS (Neuroblastoma RAS Viral (V-Ras) Oncogene Homolog), and GNL3L (Guanine Nucleotide Binding Protein-Like 3 (Nucleolar)-Like) are host dependency factors for SARS-CoV-2. To confirm this, we assessed the impact of siRNA protein knockdown on titers of SARS-CoV-2 variants Wuhan VIDO-01, and Delta B.1.617.2 (Figure 7). Knockdown of all four of these genes, KAT5, HTR3E, NRAS and GNL3L, significantly reduced the virus titers for SARS-CoV-2 Wuhan VIDO-01 and Delta B.1.617.2 variants (Figure 7), which confirmed their pro-viral role in the SARS-CoV-2 life cycle and further corroborates the robustness and proficiency of our screening methods. Western blot confirms complete knockdown of KAT5 (Figure S4) and NRAS protein (Figure 8), and qPCR quantification shows 60% and 90% reduction in HTR3E and GNL3L mRNA levels, respectively (Figure S4).

**Figure 7.**
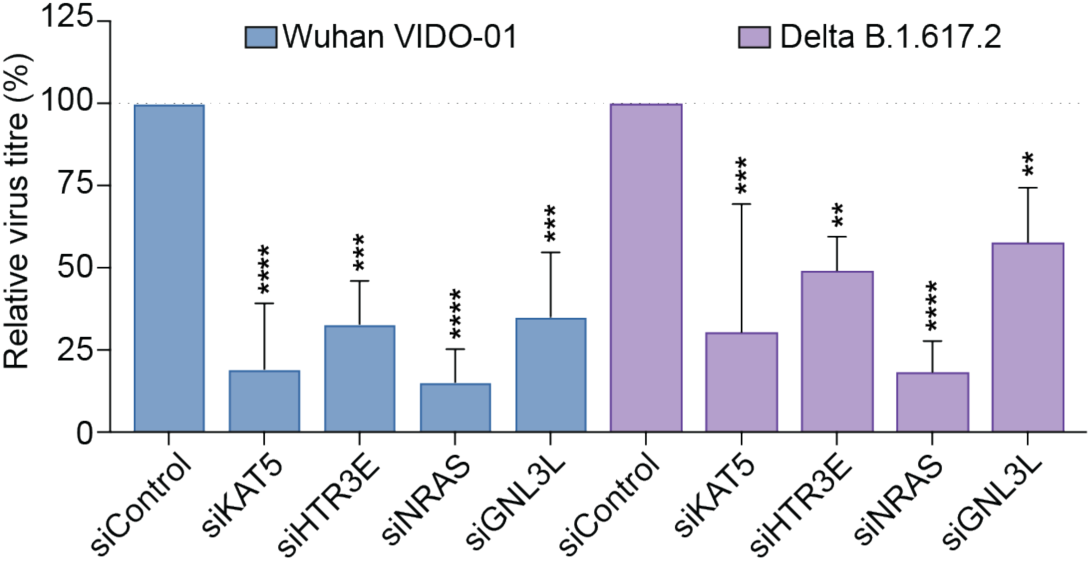
siRNA knockdown and virus titrations suggest that KAT5, HTR3E, NRAS and GNL3L genes are pro-viral for SARS-CoV-2 Wuhan and Delta variants. To confirm that siRNA knockdown of four shortlisted genes, inferred using siRNA knockdown and antiviral screenings, inhibits SARS-CoV-2 replication, NCI-H23^ACE2^ cells were transfected with pools of four siRNAs targeting each gene, followed by infection with wild-type SARS-CoV-2 Wuhan VIDO-01 (blue bars) and Delta B.1.617.2 viruses (purple bars). The x-axis indicates the siRNAs against different genes, and the y-axis denotes relative virus titers (in %), normalized to infection with a non-targeting siRNA, siControl. The data represent an average of at least three independent experiments and error bars represent standard deviation. Statistical significance was determined using two-way ANOVA and compared to siControl for each virus respectively; where ns p>0.1234, * p<0.0332, ** p<0.0021, *** p<0.0002, **** p<0.0001. [KAT5-Lysine acetyltransferase 5; HTR3E-5-Hydroxytryptamine (Serotonin) Receptor 3 family member E; NRAS-Neuroblastoma RAS Viral (V-Ras) Oncogene Homolog; GNL3L-Guanine Nucleotide Binding Protein-Like 3 (Nucleolar)-Like].

**Figure 8.**
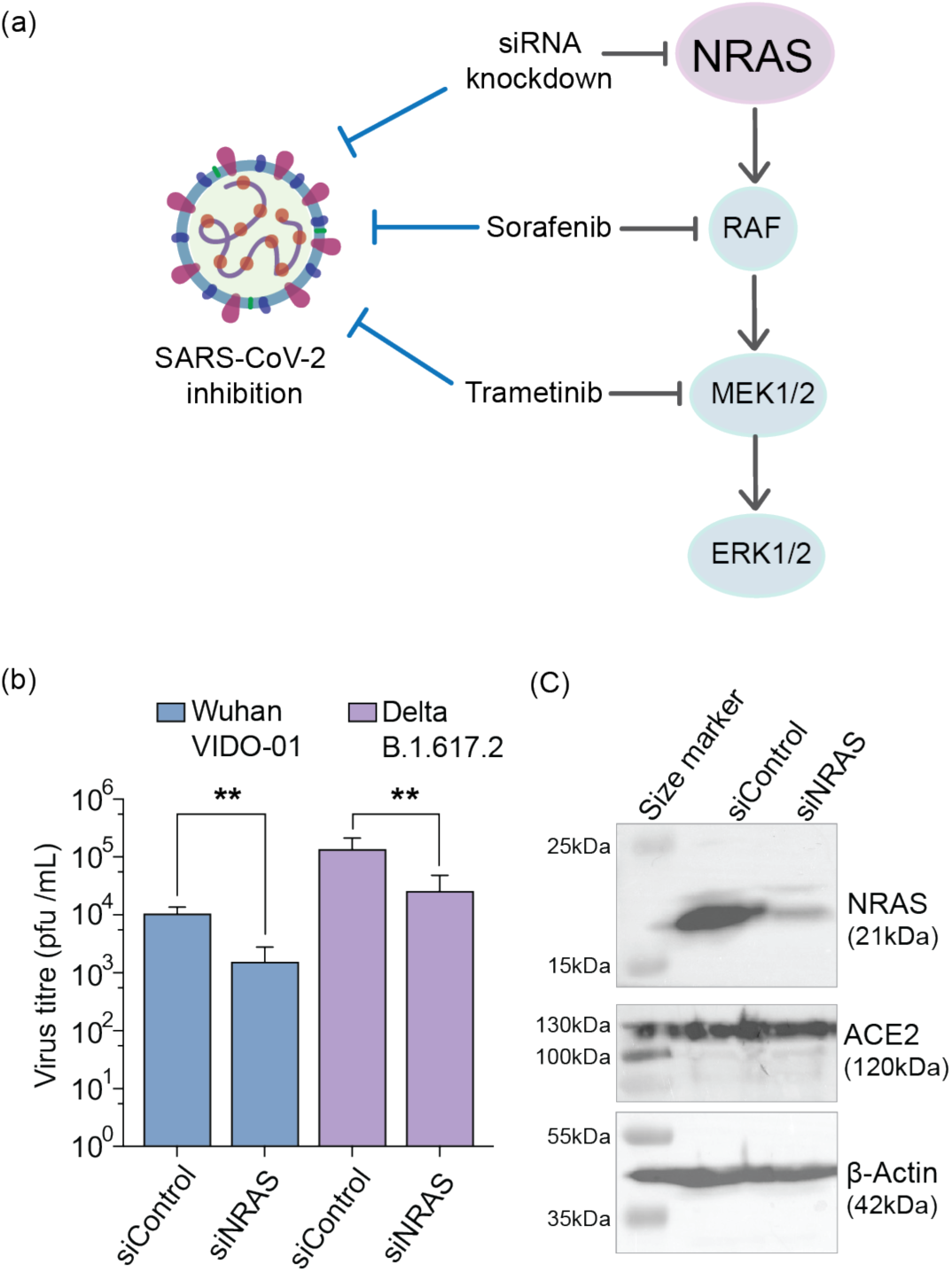
The NRAS/Raf/MEK/ERK pathway is important for SARS-CoV-2 replication. (a) siRNA knockdown of NRAS, sorafenib treatment against Raf and ERK, and trametinib treatment against MEK1/2 inhibit SARS-CoV-2, indicating that SARS-CoV-2 uses this pathway during its life cycle. [NRAS-Neuroblastoma RAS Viral (V-Ras) Oncogene Homolog; Raf-Serine/threonine kinase, rapidly accelerated fibrosarcoma; MEK1/2-MAPK/ERK Kinase or Mitogen-Activated Protein Kinase Kinase 1/2; ERK1/2-extracellular signal-regulated kinase 1/2 or Mitogen-Activated Protein Kinase 3/1]. (b) Knockdown of NRAS in NCI-H23^ACE2^ cells decrease SARS-CoV-2 Wuhan VIDO-01 (blue bars) and Delta B.1.617.2 (purple bars) virus titers significantly, with the y-axis denoting virus titers (in pfu/mL). The x-axis indicates the respective siRNA used. The data represent an average of at least three independent experiments and error bars represent the standard deviation. Statistical significance was determined using two-way ANOVA and compared to titers with a non-targeting siRNA, siControl; where ns p>0.1234, * p<0.0332, ** p<0.0021, *** p<0.0002, **** p<0.0001. (C) Western blot analysis confirmed the knockdown of protein expression of NRAS (21kDa) by ∼91.52%, normalized to expression of β-actin (42kDa). ACE2 protein expression was confirmed to remain unchanged post siRNA knockdown confirming ACE2 independent host-dependency of NRAS by SARS-CoV-2. The figure is representative of three independent experiments.

### siRNA knockdowns and drug targeting implicate the NRAS/Raf/MEK/ERK signalling pathway as pro-viral for SARS-CoV-2

The NRAS/Raf/MEK/ERK signalling pathway was identified to be required by SARS-CoV-2 in our CRISPR, siRNA knockdown, and drug screens. NRAS was identified first in CRISPR screens of HEK293T^ACE2^ and NCI-H23^ACE2^ (Figure 1f), and then, in the siRNA knockdown screen (Figure 6a, 8). MEK1/2 is targeted by the SARS-CoV-2 inhibitor trametinib (Figure 2d, f) and sorafenib targets Raf and ERK (Figure 2e, f) (summarized in Figure 8a). To validate the role of this pathway as pro-viral for SARS-CoV-2, we used siRNA knockdown of NRAS and assessed virus replication (Figure 7, 8b), followed by Western blots to confirm knockdown. Knockdown of NRAS by ∼91.52% (Figure 8c), resulted in a significant reduction of SARS-CoV-2 Wuhan VIDO-01 and Delta B.1.617.2 virus titers by up to one log-fold (Figure 8b). Furthermore, ACE2 expression was observed to be unaffected by NRAS knockdown (Figure 8c), indicating that SARS-CoV-2 hijacks the NRAS pathway in an ACE2-independent manner.

## Discussion

Although we are currently in the post-pandemic period, with fewer severe SARS-CoV-2 variants in circulation, there remains much to understand about the biology of SARS-CoV-2, its critical dependence on host cellular functions and pathways, and how we can use this information to develop antiviral therapies for SARS-CoV-2 or potentially the next coronavirus to emerge. To this end, we performed a genome-wide CRISPR KO screen with SARS-CoV-2 in a human lung cell line, that can support robust virus replication. Our study is unique in using a human lung adenocarcinoma cell line, NCI-H23^ACE2^ in which SARS-CoV-2 infection causes very high levels of cell death (up to ∼99%) post-infection. This allowed us to screen with stringent live-dead selection [30]. Our screen identified 430 enriched gene hits from NCI-H23^ACE2^ cells, and 296 enriched genes from the HEK293T^ACE2^ screen with an overlap of 18 genes identified in both screens (Figure 1a, c, e). The GO enrichment analysis of both the gene lists, implicated several cellular functions in supporting the SARS-CoV-2 lifecycle, including intracellular signal transduction, phosphorylation, intracellular transport, cytoskeleton organization, vesicle membrane function, and endosome membrane functions (Figure 1b, d).

In addition to meta-analysis, we used 2 approaches to validate the pro-viral host factors identified in our CRISPR screen-first, by testing drugs that target proteins or pathways from genes identified for ones that inhibit the virus, and secondly, by siRNA knockdown of top scoring hits and selected genes of interest. We made use of a previously developed reporter SARS-CoV-2 NLuc virus to simplify the screens [30]. Moreover, this approach also identified drugs that can be repurposed for COVID-19. The siRNA secondary screen confirmed 55 gene hits that reduced virus replication in NCI-H23^ACE2^ cells (Figure 6a). STRING network analysis on the top 55 genes that significantly reduced virus replication, provided a protein-protein interaction map of genes involved in similar functions, that may be required during SARS-CoV-2 infection (Figure 6b).

RNP complexes were one of the most enriched gene sets (GNL3L, RPL18A, RPL3, APOBEC3F, HNRNPA1, SFPQ, DDX6, and DHX35) implying the complexity and importance of the viral RNA-host protein interactions during replication of SARS-CoV-2. However, knocking down ribosomal proteins may affect the cellular translational landscape and alter normal cell survival as well as other protein interactions with SARS-CoV-2. Consistent with this, some of the siRNA knockdowns were consequently seen to reduce cell viability along with virus replication (Figure 6a). SARS-CoV-2 is, however, known to cause translational shut-off during infection [42] and may hijack ribosomal proteins as one of its mechanisms to do so. We further compared the hits corresponding to proteins associated with the RNP complexes in our siRNA knockdown screen with viral RNA-host interactome studies, and identified RPL3 [43, 44], RPL18A [43, 44], SFPQ [44], HNRNPA1 [43–45], DDX6 [45] and APOBEC3F [45, 46] to be consistently interacting with SARS-CoV-2 viral RNA. Thus, our study provides a functional confirmation to the viral RNA-host RNP complex interaction studies.

Cytokine signalling is another important pathway that was identified in the STRING analysis implicating genes such as IL36A, TRIM62, IRAK1, IFNL1, POM121, GRB2, NRAS, PSMA2, PSMB2. Severe COVID-19 in several patients has been characterized by an inflammatory cytokine storm wherein massive amounts of inflammatory cytokines are rapidly secreted in response to an infection [8]. Studies have shown that SARS-CoV-2 infection in lung epithelial cells induces transcriptional activation of inflammatory cytokine pathways and mRNA transcript levels of IFNL1 (Interferon Lambda 1), a gene hit identified in our secondary siRNA screen as being a host dependency factor (Figure 6a), was upregulated in SARS-CoV-2 infected Calu3 cells [47]. Identification of immune and inflammatory cytokines as host dependency factors is counterintuitive but since our screen relies on cell killing these cytokines may be involved in viral-induced CPE. In addition, we recently determined that the NCI-H23^ACE2^ cells used in our screens are deficient in innate immune responses, which could influence our results. Confirmation of the dependency of these factors in will require their confirmation in innate immune competent cells.

When comparing our CRISPR screen hits to the screens by other groups there are surprisingly few genes that overlap. CRISPR screen results variability between cell lines is expected but they often vary even when done in the same cell line. This could be caused by different passage of the cell lines, virus strains or CRISPR gRNA library used, MOI, and stringency of the screen indicated by the timing number of resistant cells harvested post infection. [48]. While there is little overlap between ours and others’ screens, we did see overlap of 12 and 13 genes identified between our screens in NCI-H23^ACE2^, HEK293T^ACE2^ and other CRISPR screens [21–29] (Figure 1f, S1) that highlight the importance of a few cellular pathways and protein complexes [21, 22, 24–26]. Our screen and three others identified members of the exocyst complex as proviral for SARS-CoV-2 [21, 22, 26].

Apart from important pathways, our screen also facilitated the identification of drugs that can be repurposed for COVID-19 while providing an additional means to validate the gene targets identified in the screen. Performing an antiviral screen with 21 known drugs identified two drugs that have been previously untested against SARS-CoV-2 as well as HCoV-229E and HCoV-OC43. Donepezil and dH-ergocristine are FDA-approved for cognitive disorders and in our study were found to target pro-viral genes KAT5 and HTR3E respectively [33–35]. Both these drugs effectively inhibit SARS-CoV-2 variants, Wuhan VIDO-01, Delta B.1.617.2 and Omicron BA.1 and human coronaviruses HCoV-229E and HCoV-OC43 at concentrations non-toxic to the cells (Figure 2f, 3). Donepezil is an inhibitor of acetylcholinesterase and is also shown to decrease the levels of intracellular amyloid precursor protein (APP). It does so by inhibiting endocytosis of APP, which leads to increased APP expression on cell membranes, where APP is cleaved by α-secretase enzymes [33, 49, 50]. Donepezil inhibition of KAT5 is expected to be through its inhibition of APP. APP, along with the adaptor protein Fe65, is required for transactivation of KAT5, and it stabilizes KAT5 by facilitating its phosphorylation by cyclin-dependent kinases. It is further reported to facilitate KAT5 transcription activity due to its phosphorylation activation [51, 52]. dH-ergocristine is an FDA-approved drug used for the treatment of cognitive disorders such as dementia and is a serotonin receptor antagonist [34, 35]. In our study it was found to be effective against SARS-CoV-2 variants in both, A549^ACE2^ and Calu3 cells. In Alzheimer’s studies, dH-ergocristine also acts as a direct inhibitor of γ-secretase and reduced the amyloid-β peptide [34] and SARS-CoV-2 spike protein may modulate γ-secretase activity and may affect COVID acute neuropathy [53]. In addition, several reports have suggested a link between serotonin, the immune system, and long COVID [54–56]. Thus, the pathways altered by donepezil and dH-ergocristine that inhibit virus replication may inform treatment options for the effects of COVID-19 disease on the brain.

In addition to an antiviral effect by donepezil and dH-ergocristine, siRNA knockdown of their targets, both KAT5 and HTR3E, was further observed to reduce virus titers significantly, implicating their direct role in SARS-CoV-2 replication (Figure 7). A recent study showed KAT5 regulates ZIKV replication by mediating acetylation of its NS3 helicase and the pro-viral role of KAT5 is conserved in flaviviruses including Dengue virus, West Nile virus and yellow fever virus [57]. This may allude to a similar role of acetylation by KAT5 in SARS-CoV-2 replication. This may also suggest conservation of host dependency on this protein across multiple RNA virus families. Additionally, since donepezil also inhibits the enzyme acetylcholinesterase, studying the role of this enzyme on SARS-CoV-2 replication can further elucidate the mechanism of inhibition of SARS-CoV-2 by donepezil. That donepezil as well as dH-ergocristine inhibited three coronavirus family members, also indicate conserved pan-coronavirus host dependent tendencies.

Trametinib and sorafenib, two other drugs identified in our antiviral screen can inhibit SARS-CoV-2 independently (Figure 2, 4, 5). Both the drugs modulate the NRAS/Raf/MEK/ERK pathway by inhibiting MEK1/2 and Raf respectively (Figure 8) [36, 37]. This suggests that the NRAS/Raf/MEK/ERK pathway supports virus replication. In addition, siRNA knockdown of NRAS also inhibited SARS-CoV-2 replication, thus implicating the NRAS/Raf/MEK/ERK pathway as a host dependency factor (Figure 8). The Ras/Raf/MEK/ERK signaling pathway is in the MAPK cascades and plays an important role in cell growth and proliferation. The pathway is initiated by various stimuli that can activate G protein-coupled or receptor tyrosine kinase (RTK) receptors, that in turn activate the GTPase Ras (including NRAS, HRAS and KRAS). This further activates the serine/threonine kinase Raf which promotes the kinase activity of MEK1/2, consequently activating ERK1/2 (Figure 8a). Activated ERK1/2 is responsible for phosphorylation of several transcription factors that ultimately regulate gene expression [58, 59]. Due to its important role in transcription and cell cycle regulation, the MAPK cascade is also required by other viruses including flaviviruses, enterovirus, alphaviruses, and human immunodeficiency virus (HIV), either acting as pro-viral or antiviral host factors [59–62]. An activated Ras pathway in the host cell also sensitizes the cells to reovirus infection and promotes virus spread through inhibition of IFN-β production through the RIG-I (retinoic acid-inducible gene I) signaling pathway [63, 64]. It has also been reported that inhibition of MEK1 by small molecular inhibitors, augments type 1 IFN response in the context of another respiratory virus, human rhinovirus type 2 (RV2) infections [65]. In coronaviruses, inhibition of the Raf/MEK/ERK pathway was previously confirmed to inhibit mouse hepatitis virus (MHV), a murine coronavirus, by modulating MHV RNA synthesis [66]. We speculate that the Ras/Raf/MEK/ERK signaling pathway is pro-viral for SARS-CoV-2, by inhibiting IFN-β responses and suggest that inhibition of this pathway can provide new avenues for host-directed antivirals [64, 65]. Several MAPK related biomarkers, including C-Raf, HRAS, and ERK2 were also found upregulated in PBMCs of COVID-19 patients [67]. Thus, while this pathway has been speculated to modulate replication of SARS-CoV-2, our study provides a direct *in vitro* confirmation of its involvement.

The Raf/MEK/ERK pathway may also be a pan-coronavirus modulator. Previous studies found that both, trametinib and sorafenib inhibit the SARS-CoV-2 predecessor viruses, SARS-CoV and MERS-CoV [39, 68–70] and novel to our study, we have confirmed replication inhibition by these drugs of multiple variants of SARS-CoV-2 (Wuhan VIDO-01, Delta B.1.617.2 and Omicron BA.1) as well as human coronaviruses 229E and OC43 (Figure 2, 3), suggesting that NRAS/Raf/MEK/ERK pathways may be required for all coronaviruses, and that inhibitor drugs may have anti-pan-coronavirus activity. However, that sorafenib did not inhibit HCoV-OC43 (Figure 3h) may suggest evolutionary divergence in the host requirement between HCoV-OC43 and SARS-CoV-2. Additionally, although trametinib was inhibitory to SARS-CoV-2 in both A549^ACE2^ and Calu3 cells, sorafenib did not inhibit SARS-CoV-2 in Calu3. This can indicate either a weak inhibition by sorafenib on the NRAS pathway in Calu3 cells or a possible direct targeting of viral activity by trametinib. Furthermore, trametinib or sorafenib in combination with dH-ergocristine showed more than additive inhibition against SARS-CoV-2, suggesting that targeting two host cellular pathways can lead to stronger antiviral activity while maintaining minimal cell toxicity.

Further mechanistic studies are required to highlight the role of the NRAS/Raf/MEK/ERK pathway in each step of the virus life cycle. Studies have indicated that SARS-CoV-2 alters the phosphorylation landscape of an infected cell, as well as that SARS-CoV-2 viral proteins such as N, M, S and several non-structural proteins, have functional phosphorylation sites [71–73]. Thus, we speculate that the kinase activity of the NRAS/Raf/MEK/ERK pathway components, may contribute to phosphorylation of viral proteins during infection. ERK activation is also required for transactivation of several other transcription factors, thus altering cellular gene expression to promote cell growth and differentiation [59, 74]. SARS-CoV-2 infection has been known to alter the transcriptome of a cell and induce expression of various differentially expressed genes (DEGs) [75–77]. Thus, in an alternate mechanism, SARS-CoV-2 possibly hijacks the ERK pathway to transactivate other host dependency factors or inhibit transcription of antiviral genes In conclusion, our study has used a lung cell line untested in any other SARS-CoV-2 CRISPR screen before, NCI-H23^ACE2^, and identified previously unknown antiviral activity of FDA-approved drugs, donepezil and dH-ergocristine, that may be repurposed as broad acting antivirals for coronaviruses. We also tested the inhibitory activity of kinase inhibitors, trametinib and sorafenib, against SARS-CoV-2 variants, Wuhan VIDO-01, Delta B.1.617.2 and Omicron BA.1, and HCoVs-229E and OC43 and combinations of these drugs, specially targeting multiple cellular factors or pathways, further showed more than additive inhibition. siRNA knockdown inhibition of reporter virus activity highlighted the pro-viral activity of several important genes and further testing of virus titers post siRNA knockdown of some of these genes, suggested the importance of host dependency factors KAT5, HTR3E, NRAS and GNL3L. Finally, through drug targeting and siRNA knockdowns, the NRAS/Raf/MEK/ERK pathway, an integral part of the cellular system, was identified as an important host dependency factor for SARS-CoV-2. We propose that the NRAS/Raf/MEK/ERK pathway plays a variable and possibly a central host-dependency role for SARS-CoV-2.

## Materials and methods

### Cell lines and maintenance

All the cells were maintained at 37°C with 5% CO_2_. MRC-5 were a kind gift from Dr. Linda Chelico. MRC-5, Calu3, and Vero76 cells were cultured in Dulbecco’s modified Eagle medium (DMEM) (without Sodium pyruvate) (Sigma D5796) supplemented with 10% fetal bovine serum (FBS) (Gibco 12483020) and 1x Penicillin-Streptomycin (PenStrep) (Gibco 15140122). Huh-7 cells were cultured in DMEM (with Sodium pyruvate) (HyClone SH30243.01) supplemented with 10% FBS, 1x PenStrep and 1mM non-essential amino acids. HEK293T, NCI-H23 and A549 cells were transduced with ACE2 lentiviruses using reverse transduction, and monoclonal cell selection was done as described previously [30]. NCI-H23^ACE2^ (Clone A3) cells were cultured in Roswell Park Memorial Institute (RPMI) 1640 medium (Gibco 11875093) supplemented with 10% FBS and 1x PenStrep, and 4 μg/mL Blasticidin S HCl (Gibco R21001). A549^ACE2^ (Clone B1) were cultured in F-12K Medium (Kaighn’s Modification of Ham’s F-12 Medium) (ATCC 30-2004) supplemented with 10% FBS, 50μg/mL Gentamycin Sulfate (BioBasic BS724), and 5 μg/mL Blasticidin S HCl. HEK293T^ACE2^ (Clone A2) cells were cultured in DMEM (with Sodium pyruvate) (HyClone SH30243.01) supplemented with 10% FBS, 1x PenStrep and 5 μg/mL Blasticidin S HCl. The cryomedia used for freezing the cells contained 45% complete media, 45% FBS and 10% Dimethyl sulfoxide (DMSO) (MedChemExpress HY-N7060). To test for mycoplasma contamination in all the cell lines, MycoAlert, Mycoplasma detection kit (Lonza LT07-318) was used as per the manufacturer’s protocol.

### Virus stocks

SARS-CoV-2 virus handling and related experiments were performed in Biosafety Containment Level 3 facility (CL3) at Vaccine and Infectious Disease Organization (VIDO, SK, Canada). Vero76 cells were used to prepare SARS-CoV-2 virus working stock and determine titres with TCID_50_ as described previously [30]. The following virus wild-type stocks and strains were used throughout our study - P3 (passage #3) of SARS-CoV-2/Canada/ON/VIDO-01/2020 (Wuhan1) (NCBI accession number EPI_ISL_425177), SARS-CoV-2/India/B.1.617.2 (Delta) (NCBI accession number PX393515) and P3 of SARS-CoV-2/BA.1/Omi-1 (Omicron) (NCBI accession number PX393516). The P#1 stock of SARS-CoV-2 Wuhan NLuc reporter virus was rescued from a molecular clone as described and characterized previously [30] and was used for high-throughput screening and antiviral assays. The passage 1 (p1) stock of SARS-CoV-2 Delta NLuc reporter virus was rescued from a molecular clone as described and characterized *(Rohamare et al, manuscript in preparation)*. MRC-5 was used to grow stocks and titre p4 HCoV-OC43, and Huh-7 cells were used to grow stocks and titre p5 HCoV-229E.

### Generation of Cas9 expressing stable cell lines

Cas9 lentiviruses were generated in HEK293T cells, using a Cas9 gene containing lentivirus expression vector that also contained a hygromycin selection gene, as described previously [30, 78]. Cas9 stable cell lines were generated by lentivirus transduction in polybrene containing media [78]. After 24 hours, HEK293T^ACE2^ Cas9 and NCI-H23^ACE2^ Cas9 cells were selected using 200µg/mL hygromycin B (Gibco 10687010).

### CRISPR KO screen

The human GeCKOv2 lentiCRISPRv2 KO pooled library B (GenScript SC1777) was used in this study, which contains three gRNAs targeting each of 19,050 genes, along with 1000 non-targeting control gRNAs. HEK293T cells were transfected with the pooled plasmid library to produce lentiviruses as described previously [79]. The screen was performed in triplicate with NCI-H23^ACE2^ Cas9 cells and a single replicate in HEK293T^ACE2^ Cas9 cells. Briefly, 18 x 10^6^ cells were transduced with the CRISPR lentivirus library at a multiplicity of infection (MOI) of 0.3, representing guide RNA coverage of 300x, in medium containing polybrene. 24 hours post-transduction, the cells were selected with Puromycin at 2 or 4µg/mL for NCI-H23^ACE2^ Cas9 and HEK293T^ACE2^ Cas9 cells, respectively. After 48 hours, 12 x 10^6^ cells were collected for the *T_0_* (timepoint 0) samples. 18 x 10^6^ cells were further seeded for SARS-CoV-2 infection. After 24 hours, CRISPR library transduced-NCI-H23^ACE2^ Cas9 and HEK293T^ACE2^ Cas9 cells were infected at MOI of 0.1 or 0.3, respectively, with SARS-CoV-2/VIDO-01 (Wuhan) P#3 virus. In parallel, another set of cells were treated similarly as mock infected for control sample collection. In CRISPR library transduced-NCI-H23^ACE2^ Cas9 cells, robust virus-induced cell death (∼95%) was observed, and the cells were collected 48-72 hours post-infection. For a stringent screening, CRISPR library transduced-HEK293T^ACE2^ Cas9 cells were re-infected with SARS-CoV-2/VIDO-01 twice, and cells were collected at day 6 post-infection.

At the time of cell harvesting, the cells were washed with Dulbecco’s Phosphate Buffered Saline (DPBS) (Gibco 14190250), and genomic DNA was extracted using the QIAamp DNA Blood Maxi Kit (QIAGEN 51194) as per the manufacturer’s protocol. Illumina adapters and barcodes were added to samples by PCR as previously described [79–82]. As quality control, the PCR products were confirmed to be of the desired size and purity by electrophoresis before sequencing. Samples were sequenced on an Illumina HiSeq2500 by the Michael Smith Genome Sciences Centre, BC Cancer, Vancouver, Canada. Indexed reads were demultiplexed before analysis.

### siRNA knockdown (Secondary screen and validation)

For the secondary siRNA knockdown screening, we used the “Cherry-pick custom library” tool from Dharmacon, Horizon Discovery to order ON-TARGETplus^TM^ SMARTpool siRNAs for selected genes. Genes were chosen based on statistical significance, gene functions and their involvement in other virus lifecycles. ON-TARGETplus Non-targeting Control pool (Dharmacon D-001810-10-05) was used as siControl. For individual siRNA knockdown validations, the ON-TARGETplus^TM^ SMARTpool siRNAs used include: NRAS (L-003919-00-0005), KAT5 (L-006301-00-0005), HTR3E (L-009120-02-0005), GNL3L (L-015743-01-0005) and CTSL (L-005841-00-0005). All the siRNAs were dissolved in nuclease-free water (Invitrogen 10977015), aliquoted and stored at −80°C. The reverse transfection method was used for siRNA knockdowns in NCI-H23^ACE2^ cells using Lipofectamine™ RNAiMAX Transfection Reagent (Invitrogen 13778075) as per the manufacturers protocol. Briefly, in white 96-well plates (Corning C3610), 2pmol/ well siRNAs were dissolved in Opti-MEM (Gibco 31985088), followed by addition of 0.2-0.3µL RNAiMAX reagent dissolved in Opti-MEM, incubation at room-temperature for 10-20 min, and finally the addition of 1.8 x 10^5^ cells/well diluted in RPMI supplemented with 10% FBS. To test for gene knockdown effect on virus replication, at 48 hours post transfection, the cells were infected with SARS-CoV-2 virus (strain as mentioned in each figure and result) at MOI 0.01 at 37°C for 1 hour. The virus inoculum was then replaced with RPMI supplemented with 2% FBS and 1x Pen-Strep and incubated for 24 hours at 37°C. At 24 hours post infection, the supernatant was harvested either for TCID_50_ or for luciferase assays as described below. Cell viability due to siRNA knockdown was confirmed in uninfected plates using the Viral ToxGlo^TM^ assay (Promega G8943) according to the manufacturer’s protocol and luminescence was read on Promega^TM^ GloMax® plate reader at 5 seconds integration time.

### Computational analysis of the screens

The CRISPR KO screening data was analyzed by the MAGeCK software as described previously [83]. Meta-analysis of screens published by various groups was done using Microsoft® Excel version 16.81 [21–29]. Each screen defines the top gene hits based on their CRISPR screen analysis and scoring parameters, such as log fold change and z-score. As laid out in the respective publications, the top gene hits from each screen were chosen for the meta-analysis. For meta-analysis with our NCI-H23^ACE2^ and HEK293T^ACE2^ screens, studies with no overlapping hits are excluded. These include CRISPR KO screen in Huh7 cells by Baggen *et al* [25] and in Caco2-ACE2, and Calu3 cells by Rebendenne *et al* [27]. A complete meta-analysis of all studies is represented in Supplementary Figure S1. A secondary network analysis on the top 55 hits from the siRNA knockdown validation screen was done using the STRING web resource version 12.0 https://string-db.org/.

### Antiviral assay

Based on the gene hits in our CRISPR screen, drugs were identified on the FDA database, the CancerRX database https://www.cancerrxgene.org/) and were cross analyzed on the Human gene database web resource https://www.genecards.org/. The following drugs were ordered from MedChemExpress: as 10mM dissolved in 1mL DMSO - Salirasib (HY-14754), Dihydroergocristine (mesylate) (HY-N2319), Bortezomib (HY-10227), Apremilast (HY-12085), Bezafibrate (HY-B0637), Sorafenib (HY-10201), Cytarabine (HY-13605), AdipoRon (HY-15848), Camptothecin (HY-16560), Donepezil (Hydrochloride) (HY-B0034), Metformin (HY-B0627), Trametinib (HY-10999), Homoharringtonine (HY-14944), Miconazole (HY-B0454), Helicin (HY-N7060), Tetrabromo-2-Benzotriazole (TBB) (HY-14394), and NU9056 (HY-110127), as 10mM dissolved in 1mL nuclease-free water - Flavin adenine dinucleotide (HY-B1654), and AICAR (HY-13417); and as powdered form - L-DOPA (HY-N0304), and Glutathione (HY-D0187), which were reconstituted in nuclease-free water right before use. Remdesivir (MedChemExpress HY-104077) was reconstituted in DMSO to make the main stock. Antiviral screening was done using four concentrations per drug at 5-fold serial dilutions (as mentioned in Supplementary table S5), and drug dose-response curves were generated using 6 concentrations at 2-fold serial dilutions. SARS-CoV-2 Wuhan NLuc virus was used for initial antiviral screening, SARS-CoV-2/VIDO-01 was used to generate drug dose response curves, and SARS-CoV-2/India/B.1.617.2 (Delta) and SARS-CoV-2/BA.1/Omi-1 (Omicron) were used to test drug efficacy against variants. For the antiviral assay, A549^ACE2^ cells were seeded 24 hours before infection in 96-well cell culture plates at 1 x 10^4^ cells/ well. The following day, the drugs were serially diluted as required in F-12K media supplemented with 2% FBS, 1x PenStrep and 0.1% DMSO and incubated with cells for pre-treatment for 1 hour at 37°C. After 1 hour, respective viruses were diluted in F-12K media supplemented with 2% FBS, 1x PenStrep at an MOI of 0.01 and added to cells along with diluted drugs such that the final concentration per well remains the same throughout the assay. After incubation at 37°C for 1 hour the serially diluted drugs were added to the cells again and incubated for 48 hours. The viral supernatants were harvested and titrated using TCID_50_ assays as described below. Alternatively, for the antiviral screening with the NLuc reporter virus, NLuc assay was performed as described below and luminescence was recorded. To assess inhibition of SARS-CoV-2 with drug combinations of shortlisted drugs (Donepezil, dH-ergocristine, sorafenib and trametinib), the respective drugs were first individually diluted serially, and half the volumes of each of the mentioned drug was added to the cells for treatment as described above. To calculate synergy or additive inhibition of drug combinations, the Bliss Independence Model was used [40]. The drug combinations are synergistic or more than additive if “observed inhibition of two drugs (a+b)” was greater than the “expected inhibition of the drugs (a+b)”. The “expected inhibition of the drugs (a+b) was calculated using the formula:

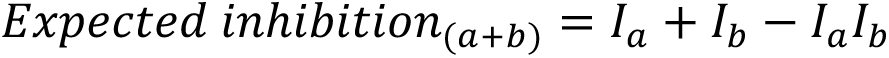

(evaluated in Supplementary table S6) [40].

To assess the cell viability under treatment, cells were treated with drugs at 37°C for 48 hours and viability assessed using the CellTiter 96® AQueous One Solution Proliferation assay (Promega G3580). Briefly, 20µL reagent was added to each well already containing 100µL media, incubated for 2 hours at 37°C and absorbance read at 490nm as an endpoint assay on the Bio-Rad xMark^TM^ Microplate Absorbance Spectrophotometer. The 50% inhibitory concentration (IC_50_) of the drug and 50% cytotoxicity concentration (CC_50_) were determined in GraphPad Prism9 using non-linear regression analysis. The selectivity index (SI) was calculated as the ratio of CC_50_/ IC_50_. The cell viability was normalized to untreated cells depicting 100% viability, whereas the virus inhibition was normalized to untreated infected cells, depicting 100% virus luciferase expression.

### Reporter luciferase assays

The Nano-Glo® luciferase assay system (Promega N1120) was used to assess NLuc expression by SARS-CoV-2 NLuc, as per the manufacturer’s protocol. Briefly, infected cells in 96-well white cell culture plates (Corning C3610) were equilibrated at room temperature for ∼5-10 min. The Nano-Glo® Luciferase Assay Reagent was prepared by combining one volume of Nano-Glo® Luciferase Assay Substrate with 50 volumes of Nano-Glo® Luciferase Assay Buffer also equilibrated to room temperature. 100µL of the reagent was added to each of the wells already containing 100µL media. The components were mixed well, incubated at room temperature for 3 minutes and Luminescence measured using the Promega^TM^ GloMax® Explorer plate reader with 5 seconds of integration time. For cell viability assays, the Viral ToxGlo^TM^ assay (Promega G8943) was used as per the manufacturer’s protocol. Briefly, 100µL of ATP detection reagent (prepared by adding ATP detection buffer to the ATP detection substrate) was added to the wells already containing 100µL media, incubated at room temperature for 10 min and the luminescence was read at 5 seconds integration time on a Promega^TM^ GloMax® Explorer plate reader.

### Virus titration using TCID_50_

Vero76 cells were seeded 24 hours prior to infection into 96 well plates, such that they were 80% confluent the next day (1 x10^4^ cells/ well). The virus to be titered was 10-fold serially diluted in deep well 96-well microplates with DMEM supplemented with 2% FBS and 1x PenStrep and then 100µL of serially diluted virus was added to the seeded Vero76 plates in 8 replicates. The plates were incubated at 37°C, and CPE was noted by microscopy at 5 days post-infection. Titers were calculated using the Spearman-Karber algorithm [84].

### Western blot

Knockdown by siRNAs was confirmed by western blot assays. Briefly, 48 hours post siRNA knockdown, cells were harvested, treated with 1x SDS lysis buffer (with 1% 1M Dithiothreitol (DTT)) and heated at 95⁰C for 10min. The proteins were then separated using a 12% SDS-PAGE gel and transferred to a methanol activated polyvinylidene difluoride (PVDF) membrane (BioRad 1620261). The membrane was blocked with 5% non-fat skimmed milk (BD Difco 232100) and probed with primary antibody overnight at 4°C followed by secondary antibody treatment at room temperature for 1 hour. The blot was developed using Clarity Western ECL substrate (BioRad 1705061) and imaged with BioRad ChemiDoc MP system. Mild-stripping was used before re-probing the blots the next day for β-actin as protein loading control (https://www.abcam.com/protocols/western-blot-membrane-stripping-for-restaining-protocol).

The antibodies used include anti-DUSP11 (ProteinTech 10204-2-AP), anti-NRAS (Abcam ab167136, 1:1000), anti-β-actin (AC-15) (Abcam ab6276, 1:10000), anti-ACE2 (R&D Systems AF933, 1:2500), anti-KAT5/Tip60 (Abcam ab151432), AffiniPure Goat Anti-Rabbit IgG (H+L) (Jackson Immuno Research 111-035-003, 1:10000), Goat anti-mouse IgG (H+L) (BioRad 1706516, 1: 10000) and Mouse anti-goat IgG-HRP (Santa Cruz sc-2354). The molecular markers used were PageRuler^TM^ Prestained protein ladder, 10 to 180kDa, (Thermo Fisher Scientific 26616) or PageRuler^TM^ Plus Prestained protein ladder, 10 to 250kDa, (Thermo Fisher Scientific 26619).

### qPCR for siRNA knockdown

To confirm siRNA knockdown of HTR3E and GNL3L, mRNA levels were assessed by qPCR. 48 hours post siRNA transfection. RNA from cells was extracted using the RNeasy kit (QIAGEN 74106) as per the manufacturer’s protocol. cDNA was prepared using the qScript cDNA SuperMix (Quantabio 95048-025) and qPCR was performed using PowerTrack^TM^ SYBR Green Master Mix (Thermo Fisher Scientific A46012) on the BioRad CFX96 Real Time System.

### Data analyses

CRISPR screening data analysis was performed using the MAGeCK software. All data barring the CRISPR screening, were analyzed and plotted in GraphPad Prism 9 software and the graphs are represented as mean +/- standard deviation unless otherwise stated. For the antiviral assays, non-linear regression model was used to generate the drug dose response curves and calculate the IC_50_ and CC_50_. The statistical analysis for each figure is indicated in the figure legends respectively. Wherever indicated, statistical significance is denoted by ^ns^ P>0.1234, * P<0.0332, ** P<0.0021, *** P<0.0002, **** P<0.0001. Western blot bands were quantified on Image Lab Software v6.1.0.

## Funding and acknowledgements

J.Q.K and J.W, conceived the research, and drafted the manuscript. J.Q.K, K.R, M.B, Y.Z, M.R, K.G, H.E, K.K.B, and J.L, carried out the experiments. J.Q.K, F.S.V and M.B created the figures, J.Q.K and F.S.V performed the computational and network analysis. All authors discussed the results and contributed to the revision of the final manuscript.

We thank Drs. Linda Chelico and Amit Gaba for providing the MRC-5 cell line, and Drs. Tom C Hobman, and Mohamed Elaish for providing the common cold coronaviruses, HCoV-229E and HCoV-43. This research was funded by a CIHR COVID-19 Rapid Research Funding Opportunity – Therapeutics Grant (VR3-172626) to J.W, F.J.V, DF. SARS-CoV-2 research in the laboratory of JW is funded by CIHR (PPE – 192112, and PPE - 190337). SARS-CoV-2 research is supported in the laboratory of D.F. by the Canadian Institutes of Health Research (CIHR; OV5-170349, VRI-173022 and VS1-175531). J.W. and D.F. are members of the CIHR-funded Coronavirus Variants Rapid Response Network (CoVaRR-Net). We gratefully acknowledge the use of infrastructure at the Phenogenomic Imaging Centre of Saskatchewan (PICS), supported by the College of Medicine, University of Saskatchewan. VIDO receives operational funding from the Government of Saskatchewan through Innovation Saskatchewan and the Ministry of Agriculture and from the Canada Foundation for Innovation through the Major Science Initiatives for its CL3 facility. J.Q.K was funded by the College of Medicine (CoM) Devolved Scholarship and Graduate Research Fellowship (GRF) from the Biochemistry, Microbiology & Immunology Department, University of Saskatchewan. M.R was funded by the College of Medicine (CoM) Devolved Scholarship and the Graduate Teaching Fellowship (GTF) from the Biochemistry, Microbiology & Immunology Department, University of Saskatchewan.

All authors declare no financial or non-financial competing interests.

## Supplementary table legends (Excel files)

**Table S1. List of gene hits from CRISPR knockout screen in NCI-H23^ACE2^ cells.** 430 gene hits were identified in NCI-H23^ACE2^ cells using MAGeCK analysis. Hits are ranked according to their enrichment score with a cut-off of 3.5 and with p-value cut-off of <0.05.

**Table S2. List of gene hits from CRISPR knockout screen in HEK293T^ACE2^ cells.** 296 gene hits were identified in HEK293T^ACE2^ cells using MAGeCK analysis. Hits are ranked according to their enrichment score with a cut-off of 3.5 and with p-value cut-off of <0.05.

**Table S3. Network analysis of 430 hits from NCI-H23^ACE2^ cells using MAGeCK software reveal enriched pro-viral pathways.** Gene ontology (GO) analysis of host-dependency hits from the NCI-H23^ACE2^ screen reveal enriched cellular networks important for SARS-CoV-2 life cycle, annotated as GOBP (gene ontology biological process), GOMF (gene ontology molecular function) and GOCC (gene ontology cellular component) as represented in Figure 1b.

**Table S4. Network analysis of 296 hits from HEK293T^ACE2^ cells using MAGeCK software reveal enriched pro-viral pathways.** Gene ontology (GO) analysis of host-dependency hits from the HEK293T^ACE2^ screen reveal enriched cellular networks important for SARS-CoV-2 life cycle, annotated as GOBP (gene ontology biological process), and GOCC (gene ontology cellular component), as represented in Figure 1d.

**Table S5. List of drugs screened in A549^ACE2^ cells against SARS-CoV-2.** 21 host-directed FDA-approved or experimental drugs were selected to be screened in A549^ACE2^ cells at 4 concentrations using SARS-CoV-2 NLuc virus. The drugs (Column A), their gene targets (identified in our CRISPR KO screen) (Column B) and concentration range (Column C) used for the screening are mentioned below. Four shortlisted drugs (donepezil, dihydroergocristine, trametinib and sorafenib) were further tested to inhibit SARS-CoV-2 Wuhan1 VIDO-01 clinical isolate, at 6 concentrations, range mentioned in the table (Column D), to generate drug-dose response curves, and one concentration each was tested against SARS-CoV-2 variants, Delta B.1.617.2 and Omicron BA.1 (Column E). The resulting inhibition of virus activity is represented in Figure 2. The four drugs (donepezil, dihydroergocristine, trametinib and sorafenib) were further tested for their efficacy against SARS-CoV-2 Wuhan NLuc and Delta NLuc viruses in Calu3 cells, at 4 concentrations, range mentioned in the table (Column G). The resulting virus activity with the drugs in Calu3 cells is represented in Figure 5. Drug combination assays against SARS-CoV-2 (wild type and NLuc viruses) was done using the calculated IC_50_ values from Figure 2, in A549^ACE2^cells (Figure 4, Column F) and Calu3 cells (Figure 5, Column H).

**Table S6. Calculation of synergistic inhibition of SARS-CoV-2 Wuhan, Delta and Omicron variants, by combination of drugs in A549^ACE2^ and Calu3 cells using the Bliss Independence Model.**

**Table S7. siRNA knockdown screen in NCI-H23^ACE2^ cells with SARS-CoV-2 NLuc virus.** 165 genes were selected to be validated from the CRISPR KO screens by siRNA knockdowns. Hits are ranked by percent luciferase activity (lowest to highest) assessed 24 hours post infection normalized to siControl, with +/- standard deviation (SD) mentioned against each gene, along with the corresponding percent cell viability normalized to siControl in uninfected cells with +/- SD. The top 55 genes (shaded in grey) with significant reduction in infection are represented in Figure 4.

---

**Figure S1.**
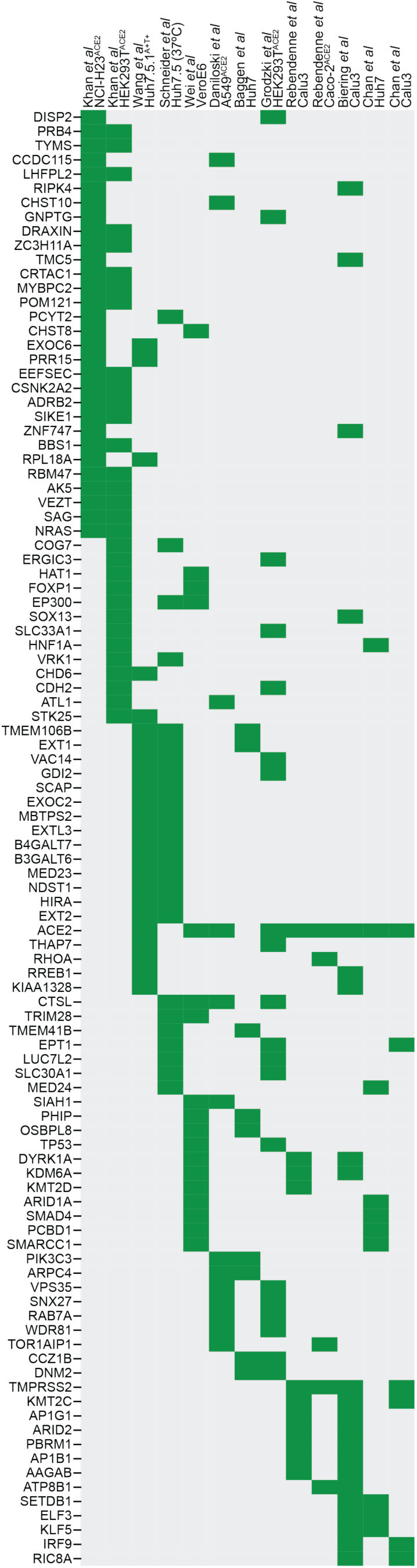
Meta-analysis of multiple CRISPR screens identify common SARS-CoV-2 host-dependency factors. A complete meta-analysis of 10 CRISPR KO studies done between 2020 to 2023, in multiple cell lines (as labelled above the graph) [21–29], reveal overlapping gene hits, represented as green blocks; with the genes labelled on the left. The genes are arranged according to the p-value calculated for NCI-H23^ACE2^ and HEK293T^ACE2^ screen in our study, followed by common clusters of genes from other groups.

**Figure S2.**
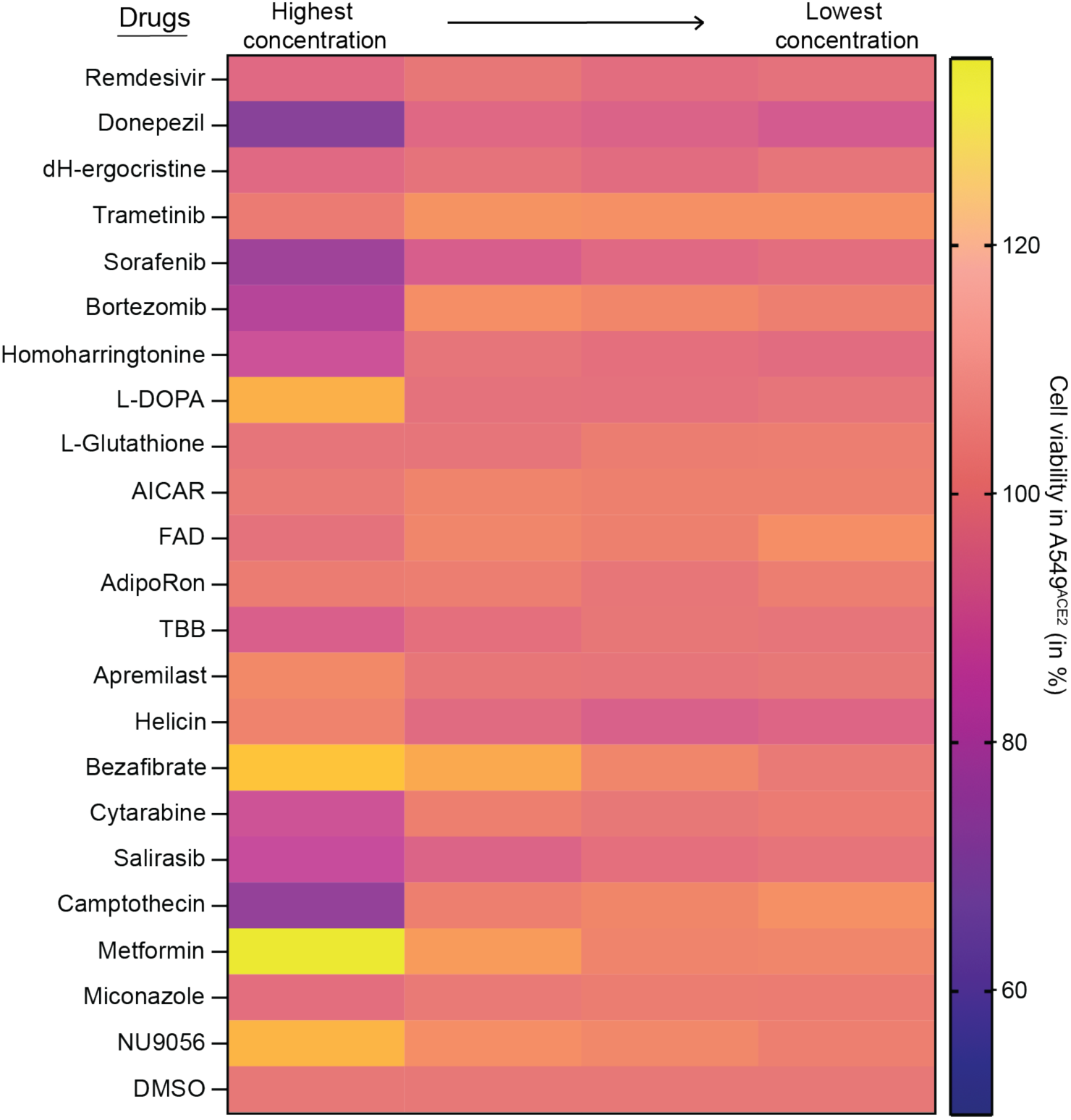
Treatment with drugs from the antiviral screen did not adversely affect cell viability in A549^ACE2^ cells. Using an ATP based cell viability assay, all the drugs tested in the antiviral screen in Figure 2a, were confirmed to not be toxic to A549^ACE2^ cells at corresponding concentrations. Cells were incubated with the drug for 48 hours before testing the cell viability and is represented in the heatmap as relative cell viability (in %), normalized to DMSO treated cells (represented at 100% as orange bars). The scale of cell viability is indicated on the right, with all drugs maintaining cell viability of >70%. The data represent an average of at least three independent experiments.

**Figure S3.**
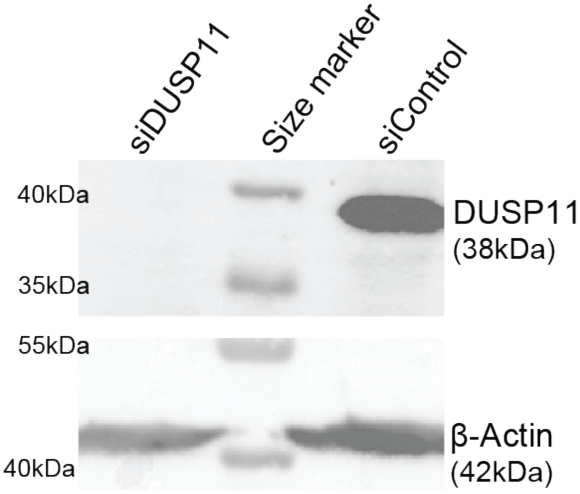
siRNA knockdown and transfection competency of NCI-H23^ACE2^ cells was confirmed by DUSP11 knockdown. Western blot analysis confirmed the knockdown of protein expression of DUSP11 (38kDa), normalized to expression of β-actin (42kDa). The figure is representative of three independent experiments.

**Figure S4.**
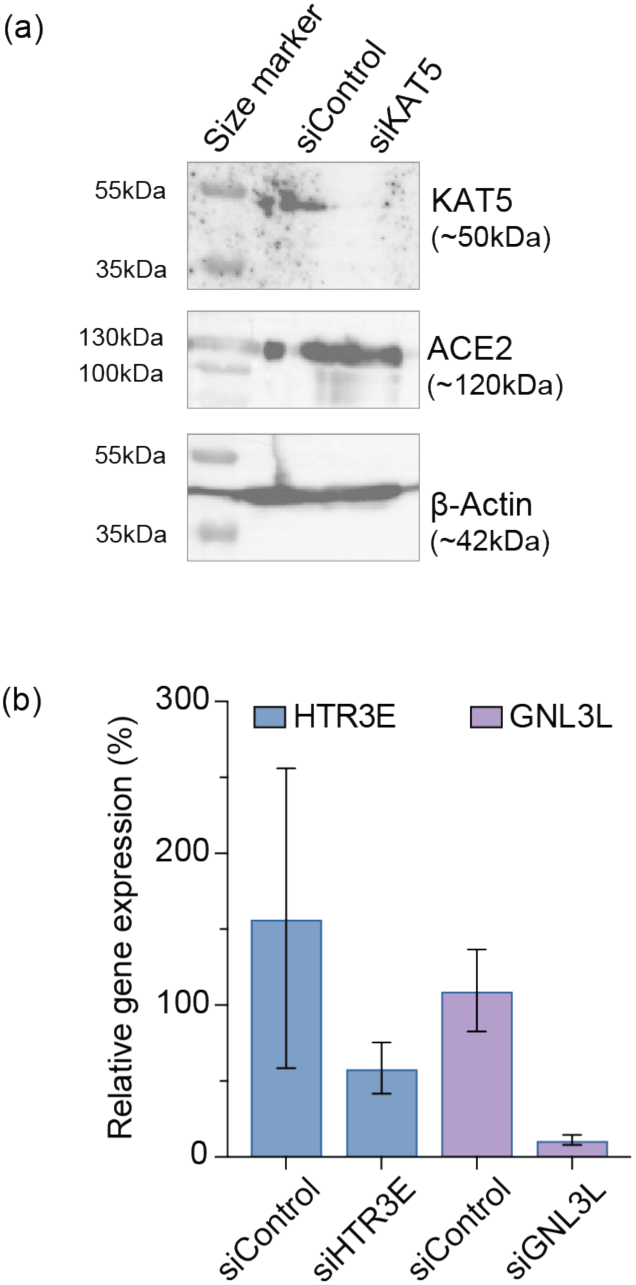
Validation of siRNA knockdown in NCI-H23^ACE2^ cells was performed by Western blot analysis or qPCR quantification. (a) Western blot analysis confirmed complete knockdown of protein expression of KAT5 (50kDa), normal to expression of β-actin (42kDa). ACE2 protein expression was confirmed to remain unchanged post siRNA knockdown. (b) qPCR quantification confirmed the siRNA knockdown by showing reduction in mRNA expression of HTR3E by about 60% (blue bars) and GNl3L by about 90% (purple bars), as compared to mRNA expression in cells targeted by a non-targeting siRNA, siControl.

